# Multi-Objective Bayesian Optimization for Data-Efficient Bioprocess Development

**DOI:** 10.64898/2026.02.02.703372

**Authors:** Edward Ma, James Morrissey, Shutong Duan, Ziqi Lu, Sandeep Ranpura, Sneha Arora, Anita Dabek, Chen Liu, Gheorghe Alexandra-Gabriela, Louis K. W. Fong, Maria Sani, Pavle Vrljicak, Deniz Demirhan, Michael Betenbaugh

**Affiliations:** Department of Chemical and Biomolecular Engineering, Johns Hopkins University, Baltimore, USA; Lonza Biologics, Slough, United Kingdom

## Abstract

Process optimization for Chinese hamster ovary (CHO) cell culture remains a challenge in biopharmaceutical development because multiple interacting parameters jointly influence productivity and product quality attributes. Traditional design-of-experiments (DoE) methods, while systematic, become impractically expensive when extended across multiple parameters and clones. To address this challenge, we developed a multi-objective Bayesian Optimization (BO) framework that identifies optimal process conditions efficiently in grouped recommendations, which is well suited for experimental workflows in bioprocess development. The model integrates continuous variables such as pH, DO, temperature, and feed rate with categorical identifiers to enable knowledge transfer across clones and scales, optimizing titer, glycan profile, and charge variants. We validated the framework through in-silico benchmarks on analytic functions, retrospective cross-validation on historical CHO datasets, and forward experimental validation in small-scale bioreactors. Across these tests, our algorithm consistently outperformed Latin Hypercube Sampling (LHS) and Random Search baselines, achieving superior performance under a limited experimental budget. The framework improved titer by up to 37% under single-objective optimization. In the multi-objective setting, it increased titer by 25% while simultaneously reducing overall glycan-profile error by a factor of seven, demonstrating the ability to optimize multiple biologically coupled objectives simultaneously. Through comprehensive in-silico and experimental validation, this study establishes a framework that enables adaptive, AI-guided process development and improves decision-making across multiple objectives, clones, and scales while minimizing experimental runs in process development and optimization workflows.

## 1. Introduction

Chinese Hamster Ovary (CHO) cells are the predominant mammalian host for the production of recombinant therapeutic proteins, particularly monoclonal antibodies^1–4^ due to their stable and adaptable growth, ability to perform human-like post-translational modifications, and established regulatory acceptance^5,6^. Fed-batch culture is the industry standard for CHO-based biomanufacturing, offering high cell densities and product titers through controlled nutrient addition and process parameter modulation. However, the complexity of CHO cell metabolism, combined with many confounding and dynamic variables in fed-batch culture, poses significant challenges for process understanding, optimization, and control^7^. Variability in cell growth, nutrient utilization, metabolite accumulation, and critical quality attributes (CQAs) such as glycosylation arise from intricate interactions between process parameters and cellular physiology^8–10^. Beyond these inherent metabolic and process-driven factors, clone-to-clone variability, arising from differences in transgene integration, expression capacity, epigenetic state, stress responses, and metabolic flexibility, introduces an additional layer of uncertainty that complicates process predictability^11–13^. This challenge is further amplified by the rapid expansion of new product modalities, such as bispecific antibodies and Fc-engineered or heavily glyco-engineered molecules, which impose distinct metabolic burdens and quality-attribute requirements, often altering process sensitivity and increasing the divergence between optimal operating conditions across cell lines^14–16^. Accordingly, the present study includes data from CHO clones producing both a monoclonal antibody and a bispecific antibody to evaluate optimization performance across both clonal variation and product modality.

In an ideal world, a full-factorial Design of Experiments (DoE) would map main effects and interactions across the tested variable domains and localize the optimum and have historically served as the foundation for process optimization^17,18^. However, in practice, the experiment count explodes exponentially for implementation of a full DOE. A three-level designs require 3^n^ experiments for n variables, and the burden multiplies when individual clones must be tuned, making such designs impractical under typical timeline and resource constraints. Industrial standards starting on a new production platform now default to lighter designs, such as fractional factorials, Box-Behnken, or central-composite response-surface methodologies (RSM), that capture the dominant effects with far fewer runs^19^. For example, a recent CHO cell study optimized trace-metal supplementation with a three-level DoE implemented in Sartorius MODDE Pro^20^. In parallel, ‘Space-filling’ designs, especially Latin-hypercube sampling (LHS), are widely used to generate new data points that effectively spread across entire variable domains when historical data exists but not suffice for DoE and RSM construction^21^. An example is a 2024 study that drew 40 LHS samples, fitted an RSM, and then solved for the optimum directly on the fitted surface^22^. While these new industrial standard designs are effective, they sometimes miss the optimal operating conditions and are still highly resource intensive. Alternatively, Bayesian Optimization (BO) provides an adaptive, data-efficient option for streamlining bioprocess design for enhanced process performance^23^.

In this study, we developed and applied a BO framework to CHO cell fed-batch processes for recombinant protein production. By combining probabilistic surrogate modelling with acquisition-driven experiment selection, BO iteratively balances exploration and exploitation to identify improved process operating conditions under tight experimental timelines and budgets^24,25^. Moreover, Bayesian inference facilitates adaptive learning as new data become available. This makes it particularly well suited for experimentation in settings where each trial is expensive or slow to perform, such as in chemical and biological process development^26–29^. In prior work, we developed a multi-recommendation BO that delivered consistently better optima than classical RSM strategies for a 3-variable basal-media design to optimize titer for the same experimental budget^30^. However, bioprocess optimization is more than just maximizing titer, as other product qualities such as charge variants and glycan profile also contribute to process performance^16,31,32^. Moreover, practical optimization often spans multiple clones and culture scales, requiring a framework capable of handling both dynamic process parameters, clonal differences, and reactor scale.

The present work extends on our prior framework to a BO algorithm capable of simultaneously optimizing multiple objectives across multiple CHO clones and scales. The framework identifies the optimal continuous process parameters (e.g., pH, DO, temperature, and feed rate) at a desired categorical state (e.g., clone, bioreactor scale) that optimizes product titer, glycan profiles and charge variants. Using experimental data from small-scale bioreactors (ambr®15), we demonstrate the application of Bayesian methods for process calibration and prediction. The method integrates Gaussian Process (GP) surrogates with an Tchebycheff scalarization strategy to navigate trade-offs between potentially competing objectives. In addition, the algorithm incorporates a multi-recommendation batch selection mechanism based on k-medoids clustering.^33^ At each iteration, the optimizer draws a large set of candidates and groups them into different clusters. The most representative and centrally located point of each cluster is chosen as part of the batch recommendation. This strategy ensures that the recommended conditions are both potentially valuable and diverse, enabling parallel experiments that explore multiple distinct regions of the design space rather than repeatedly sampling a narrow local optimum.

Alongside predictive accuracy, probabilistic optimization must also be validated on how well its uncertainty is calibrated and how reliably that uncertainty improves decision-making. To address this, we evaluated our approach in two complementary ways. We first benchmarked against LHS and Random Search across analytic test functions and biologic surrogates to assess convergence behaviour and objective minimization of our BO platform because LHS and Random Search represent established, model-free baselines commonly used in bioprocess exploration and DOE-style screening^21,34^. These methods provide a neutral point of comparison for measuring whether a model-based optimizer such as BO can outperform unbiased, uniformly sampled exploration. Second, we conducted offline replay with a random-forest (RF) ‘teacher’ trained on historical data from multiple clones and scales to test more relevant solution space landscape and generalizability. Finally, we conducted new fed-batch experiments to validate predictive capability of experimental performance. Together, these checks assess prediction, calibration, and policy quality and increase confidence that observed improvements reflect genuine progress toward the global optimum. This approach provides a foundation for enhancing process understanding and enabling model-based optimization strategies for next generation CHO cell bioprocesses.

Taking together, these gaps highlight the need for a unified, data-efficient optimization framework that can adapt across clones, scales, and competing performance objectives. Although BO has demonstrated promise in small-scale or single-objective settings, its application to realistic CHO cell bioprocesses, where categorical factors, nonlinear responses, and clonal heterogeneity interact, remains largely unexplored. Here, we address this need by applying our multi-recommendation Bayesian framework to handle multi-objective optimization across continuous process parameters, such as pH and temperature, and categorical bioprocess states, such as clones and scales. We demonstrate that this approach not only accelerates the search for improved operating conditions but also provides a principled mechanism for navigating trade-offs between productivity and product quality. By combining analytic benchmarking, offline historical training, and forward validation experiments, this study establishes a practical and generalizable foundation for model-based CHO cell process development. Our findings illustrate how probabilistic, uncertainty-aware optimization can complement and enhance traditional DoE and RSM approaches, ultimately enabling more efficient, informed, and scalable decision-making in next-generation biomanufacturing.

## 2 Methods

### 2.1 Bayesian model development

#### Model Structure

A Gaussian Process (GP)-based Bayesian Optimization (BO) framework was developed for multi-objective bioprocess parameter optimization. The model uses a Matern kernel to capture nonlinear relationships between process variables and target responses^30,35^. The framework accommodates both continuous process variables (e.g., pH, Temperature, Dissolved oxygen (DO), and feed rates) and categorical variables (e.g., clone identity, culture scale), enabling simultaneous optimization and transfer learning of patterns across different platforms and scales. An overall algorithm scheme is illustrated in Figure 1. The process begins with preprocessing and encoding of historical observations, followed by hyperparameter (e.g., length scale) sampling and GP model fitting for each objective. The algorithm then constructs multi-objective acquisition surfaces, aggregates candidate optima, and applies K-medoids clustering to identify a diverse batch of recommended operating conditions. These selected points are evaluated to generate multi-objective predictions and are applied for experimental validations.

**Figure 1.**
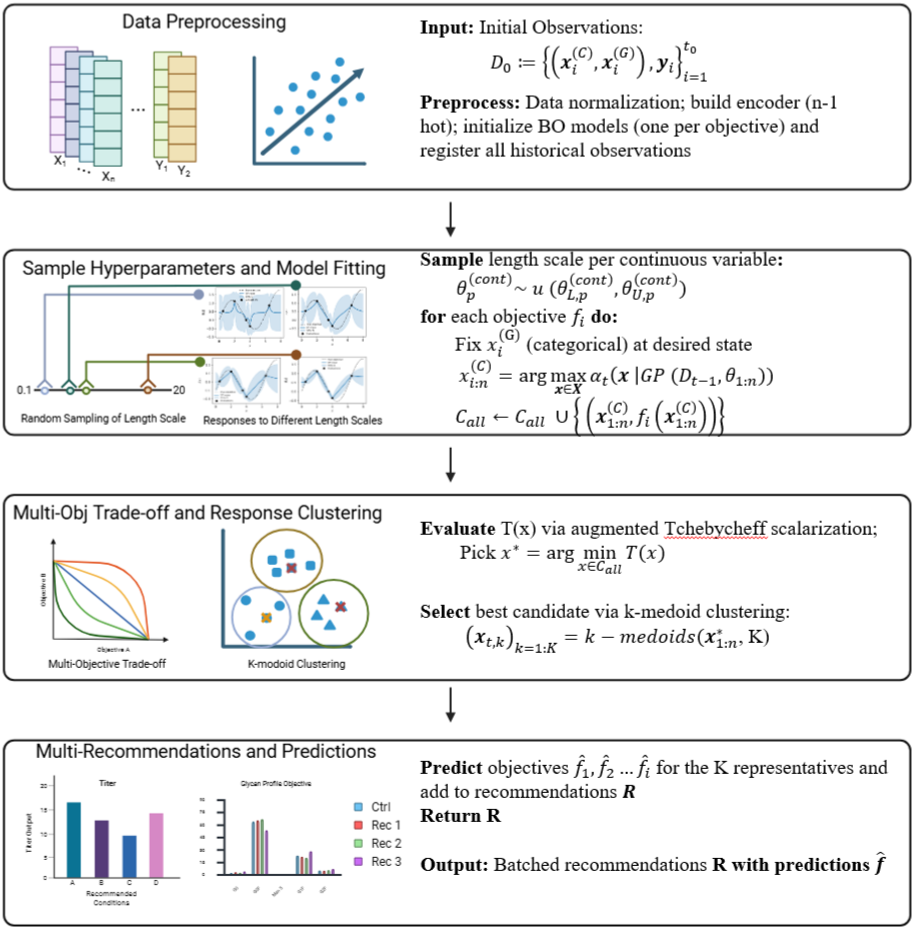
Workflow of the K-size Batched Multi-Objective Bayesian Optimization algorithm. The schematic illustrates four major stages of the proposed batched optimization framework: (1) data preprocessing and model initialization, (2) random sampling of Gaussian process hyperparameters and model fitting, (3) multi-objective acquisition evaluation with Tchebycheff scalarization followed by response-space clustering, and (4) generation of K batched recommendations with predicted outcomes for experimental testing. The diagram corresponds to the detailed steps given in Algorithm 1.

#### Continuous and Categorical Variables

Variables were grouped as continuous (pH, dissolved oxygen, feed rates, temperature) or categorical (clone identity and scale). The categorical factors were encoded using an (n-1)-hot encoding strategy^36^. Each categorical variable with n possible classes was represented by n – 1 binary indicators. For example, with three clones, clone A was encoded as [0,0], clone B as [1,0], and clone C as [0,1]. This approach removes redundant degrees of freedom in categorical representation, preventing collinearity among variables while allowing the model to incorporate categorical information numerically without introducing bias. Although Gaussian process surrogates do not inherently model categorical variables, clone identity and scale are treated as fixed contextual identifiers during optimization. This constrains all proposed solutions to valid experimental categories while allowing the surrogate model to leverage shared structure across contexts, making the one-hot encoding strategy appropriate in this setting.

#### Acquisition and Batch Recommendations Strategy

To guide the BO predictions, historical bioprocess data on several cell lines were used to both train and validate the model. Various combinations of pH, DO, temperature, and feed rate under different culture scales were tested within a defined solution space. A mapping between these process parameters and product attributes such as titer, glycan profile, and charge variants was constructed by performing regression using GP. The prior mean and variance of the GP model was calculated as described in equation 1:

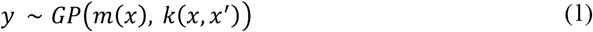

where *y* is the product attribute output, *m*(*x*) is the prior mean of input vector x, *k*(*x, x*^′^) is the covariance function.

The posterior mean *μ*(*x*_∗_ | *D*) and variance *σ*^2^(*x*_∗_ | *D*) on query point at unseen input points *x*_∗_ will then be evaluated by the following equations:

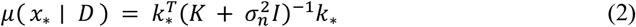

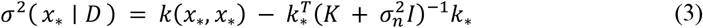

Candidate points for the next round of experiments were selected using Upper Confidence Bound (UCB), with the exploration-exploitation trade-off using the parameter *κ* as defined in equation 4:

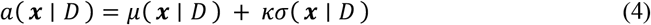

where *a*(***x*** | *D*) is the acquisition function value at any input ***x*** ∈ ℝ^*d*^, *μ*(***x*** | *D*) is the predicted mean of the surrogate GP model at input process parameter vector ***x***, *σ*(***x*** | *D*) is the predicted standard deviation, and *κ* is the exploration-exploitation trade-off parameter^37^. The first term, *μ*(***x*** | *D*), guides the search toward high performance (exploitation), while the second term, *κσ*(***x***), encourages sampling in solution space of high uncertainty (exploration).

Multi-Scale Multi Recommendation (MSMR) approaches were implemented for batch recommendations^33^. Random length scales (*ℓ*) were sampled for each input process parameter variable, *ℓ* ∈ [0,1, ^2^0] with an over-sampling rate of 20. The candidates were clustered using a k-medoid method to generate n final recommendations, where the number of recommendations per iteration is denoted as *nrec*.

In comparison as a baseline method, LHS is a model-free, space-filling experimental design method that does not rely on any model or prior knowledge of the response surface. Instead, it partitions each input variable into equally probable intervals and samples once from each interval and enforces a stratification constraint such that each interval is sampled exactly once per variable, ensuring uniform marginal coverage of the search space. By spreading samples evenly across all dimensions, LHS aims to increase the likelihood of discovering high-performing regions. As a static, unsupervised sampling strategy, it does not incorporate feedback from observed outcomes, a limitation shared by other model-free design methods. Bayesian optimization, in contrast, is inherently adaptive, using observed data to iteratively balance exploration and exploitation.

#### Multi-Objective Recommendation

To handle multiple objectives, separate Gaussian processes were trained for every objective. The top candidates were collected to form a joint pool 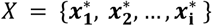, where *i* is the number of objectives. For every ***x*** ∈ *X*, we computed the posteriors *μ*_*i*_(*x*), *σ*_*i*_(*x*) and formed *i* an ideal point estimate *z*^∗^ and nadir 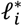 using upper and lower confidence bounds:

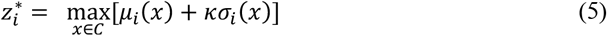

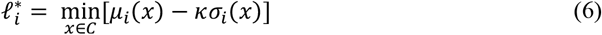

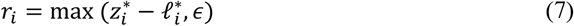

For a two-objectives problem, the candidates were then scored using the augmented, scaled Tchebycheff function:

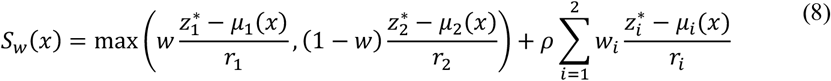

Where 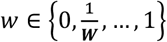 is the preference weight on the pareto front and *w*_1_ = *w, w*_2_ = 1 −*W*. For each w, a finalist candidate *x*_*w*_ = arg min *S*_*w*_(*x*) is picked^38^.

### 2.2 Model Benchmark

#### In-silico validation with empirical equation systems

BO was benchmarked sequentially against LHS and Random Search on multiple analytic/surrogate objective functions. LHS and Random Search were selected as benchmarking baselines because they represent standard non-adaptive experimental design strategies commonly used in bioprocess optimization. Both methods explore the design space without incorporating prior knowledge or iterative learning. By comparing BO against these non-adaptive approaches, we demonstrate its capabilities in efficiently balancing exploration and exploitation to identify optimal process conditions under limited experimental budgets. During benchmarking, different configurations of BO were tested. All objectives were cast on a ‘down-is-better’ scale. Minimum loss at each iteration was reported.

#### Validation on historical data (offline teacher)

The developed BO was benchmarked against LHS and Random Search for a multi-objective bioprocess optimization task. To establish a fair comparison, all methods were examined under the same experimental budget (number of experiments), with each BO run consisting of n recommendations per iteration until the budget was reached. For the benchmark, we established a teacher-student offline evaluation framework, where the ‘teacher’ model simulates the ground truth response surface and the ‘student’ model utilizes the optimization policy being tested. To evaluate the robustness of model performance, we conducted a five-fold evaluation, in which the dataset is partitioned into five equal subsets. In each round, 80% of the data (four folds) is used to train the offline teacher model, and the remaining 20% (one fold) is held out for validation. This process is repeated five times so that each subset serves as the validation fold exactly once. To account for stochastic variation such as model noise and initialization effects, we repeated this procedure using five randomly selected data partitions and three random seeds per partition, yielding 15 independent evaluations for each model. For each run, we computed the AUC as a quantitative measure of optimization efficiency, where a higher AUC reflects faster convergence towards the optimum during earlier iterations. We then calculated the change in AUC (ΔAUC) as the difference between the BO method and the Random Search for each fold–seed combination. To estimate the variability and confidence of this improvement, we performed bootstrap resampling on these 15 ΔAUC values, repeatedly sampling with replacement to generate 10,000 mean estimates. During the benchmark, we tested BO policies with different exploration-exploitation balances (k) and varying numbers of recommendations per iteration (nrec). Performance was evaluated using two primary measures: (1) the area under the convergence curve (AUC) averaged across repetitions defined as

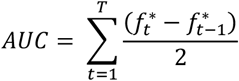

where 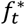 denotes the best objective value obtained up to iteration t. This metric quantifies the cumulative rate of improvement, with higher AUC indicating faster and more stable convergence toward the optimum; and (2) the mean optimality gap, defined as the difference between the best value found by the optimizer and the teacher-model optimum.

#### Validation with new experimental data

We defined five validation objectives to cover knowledge transfer, quality attributes, and multi-objective behavior. All runs used the same number of recommendation per batch (n_rec_ = 3), and exploration parameter (k = 1). Details and the rationale for all tests are provided in Table 1.

**Table 1.** Experimental Validation Design (250-mL bioreactors). All cases use the default BO configuration (k = 1; nrec = 3) and the same evaluation protocol. For each clone, we performed a control condition. Cases 1 and 2 test a single-objective on the titer with and without learning from multiple clone data. Cases 3 and 4 test optimization capability for product quality attributes. Case 5 tests multi-objective optimization (titer + glycan).

| Case # | Training Data | Objective | Target Cell line | Training Purpose |
| --- | --- | --- | --- | --- |
| 1 | Clone B | Titer | B | Within Clone single-objective (titer) Control |
| 2 | Clone A + B | Titer | B | Transfer for titer uses A+ B data |
| 3 | Clone B | Glycan Profile | B | Product Quality Attribute |
| 4 | Clone B | Charge Variant | B | Product Quality Attribute |
| 5 | Clone A | Titer/Glycan Profile | A | Multi-objective optimization (titer + glycan) |

For all clones, namely clone A (monoclonal antibody), clone B (Bispecific antibody), a control condition was run at pH 7.06, 60% DO, 33°C, and 1x feed rate to establish the baseline. The domains were: pH ∈ [6.7,7.4]; DO ∈ [20, 80]%; Temperature ∈ [28,36] °C; and feed rate ∈ [0.5, 1.5] multiplier.

For the glycan profile objective, the target composition was defined as 4.1% G0, 52.8% G0F, 1.2% Man5, 26.1% total G1F, and 7.8% G2F based on historical process data. For the charge variant objective, the target distribution was defined as 35% acidic, 50% neutral, and 15% basic species. Both optimization objectives were formulated as minimizing the mean squared error (MSE) between the predicted glycan distribution and this target setpoint, as defined in Equation:

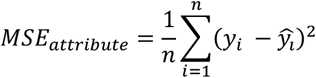

### 2.3 Experimental methods

#### 2.3.1 Historical Data (Used for Model Training and Retrospective Validation)

##### Cell lines, culture scale, and datasets

CHO cells producing therapeutic antibodies were used to generate the historical datasets summarized in **Table 2**. Clone A produces a monoclonal antibody (mAb), while Clone B produces a bispecific antibody (BiSAb). Historical data were collected across two culture scales: ambr®15 (15 mL working volume) and single-use 250 mL bioreactors. Cells were maintained in Lonza’s chemically defined, serum-free medium and expanded for 4–5 passages prior to inoculation of production cultures.

**Table 2.** Summary of the historical CHO bioprocess dataset used for model training, including clone identity, produced molecule, culture scale, and corresponding number of experimental runs.

| Clone | Molecule | Scale | Dataset Size |
| --- | --- | --- | --- |
| A | mAb | 15 mL | 36 |
| A | mAb | 250 mL | 6 |
| B | BiSAb | 250 mL | 16 |

##### Historical fed-batch process and parameter space: ambr®15 system

Historical fed-batch runs for Clone A were conducted in the Sartorius ambr®15 system following Lonza’s standard fed-batch process. Inoculum cultures were expanded from shake flasks to roller bottles at 36.5 °C, 5% CO_2_, and 140 rpm, before seeding N-stage production cultures at an initial viable cell density of 0.5 × 10^6^ cells/mL. Production cultures were grown in Lonza production medium for 15 days.

Within the ambr®15 dataset (36 runs shown in **Table 2**), process parameters were systematically varied to define the model training space. The explored parameter space included pH, temperature, DO, and feed rate, with a shift in these parameters occurring post exponential growth phase^39^. Temperature and pH shifts were applied between growth and production phases according to internal process guidelines. DO was controlled via gas sparging and agitation cascades, and feed rates were determined using a proprietary viable cell concentration–based feeding strategy. This space-filling design provided sufficient coverage of the multidimensional process space for surrogate model training and retrospective evaluation.

##### Historical fed-batch process: 250 mL bioreactor system

Additional historical fed-batch runs were conducted in single-use 250 mL bioreactors operated by Culture Bioscience for both Clone A (6 runs) and Clone B (16 runs), as summarized in Table 1. The overall process design mirrored the ambr®15 process, including inoculation strategy, feeding approach, temperature shift schedule, and DO control scheme. These datasets provided complementary information at a larger working volume and supported cross-scale and cross-clone model training.

##### Dataset characteristics

Across both scales and clones, the historical datasets exhibited substantial variation in cell growth, metabolic profiles, titer, glycan composition, and charge variant distribution. This diversity ensured adequate coverage of biologically relevant operating regimes and provided a suitable foundation for training surrogate models and offline teacher models used in retrospective benchmarking.

#### 2.3.2 Forward Experimental Validation (Used to Test BO Recommendations)

Forward experimental validation of BO recommendations was performed in single-use 250 mL bioreactors operated by Culture Bioscience. These experiments were designed to test BO-recommended operating conditions under controlled bioreactor conditions.

Five forward validation cases were conducted, as summarized in **Table 1**. Cases 1 and 2 evaluated single-objective titer optimization for Clone B, using within-clone training (Clone B only) and across-clone training (Clone A + B), respectively. Cases 3 and 4 evaluated single-objective optimization of glycan profile and charge variant distribution for Clone B. Case 5 evaluated multi-objective optimization of titer and glycan profile for Clone A using within-clone training. For each case, the BO model was trained on the specified historical dataset and used to generate a batch of recommended operating conditions, which were then implemented experimentally. The bioreactor process design followed the same inoculation strategy, feeding approach, temperature shift schedule, and dissolved oxygen control scheme used in the historical 250 mL datasets. Cultures were inoculated at 0.3–0.5 × 10^6^ cells/mL and operated under automated control of pH, DO, and temperature. Nutrient feeds and glucose were added as required to maintain concentrations within predefined limits.

##### Sampling and analytical assays

During forward validation experiments, daily samples were collected to measure viable cell density and viability using trypan blue exclusion on a Vi-CELL analyzer. Metabolites, including glucose, lactate, glutamine, and ammonia, were quantified using a Nova biochemical analyzer. Product titer was measured by Protein A HPLC, charge variants by icIEF, and glycan distributions by the Gly-X assay.

## 3 Results

### 3.1 In-silico verification

We first evaluated the BO algorithm (Methods section 2.1) against LHS and Random Search using 15 benchmark functions (Supplementary Table 1). These benchmarks include a mix of standard analytic test functions and process-inspired surrogate functions designed to reflect characteristics of real bioprocess systems. All benchmarks were formulated as minimization problems, such that lower objective values correspond to better performance.

The benchmark set spans a range of optimization difficulty, including smooth convex landscapes, highly multimodal functions with many local optima, and noisy, nonlinear response surfaces typical of bioprocesses. Across all benchmarks, BO consistently outperformed both LHS and Random Search under both exploitative (*κ* = 1) and explorative (*κ* = 5) configurations. When averaged across analytic and bioprocess surrogate benchmarks, BO improved final objective values by approximately 68.5% for the analytic functions and 18.8% for bioprocess surrogate problems relative to Random Search.

Six representative benchmarks are shown in **Supplementary Figure 1**, including three analytic functions and three bioprocess-inspired surrogates. These functions were selected to illustrate increasing optimization complexity. The analytic benchmarks **(Supplementary Figure 1a–c)** are widely used to evaluate optimization behavior. Ackley5 contains many local minima that challenge algorithms to escape premature convergence, while Hartmann6 is a smooth but strongly multimodal function used to assess convergence accuracy. In contrast, Branin is a relatively simple function with a small number of global minima, allowing most methods to perform well.

Across the analytic benchmarks, BO achieved faster convergence and lower objective values than both LHS and Random Search. For simpler landscapes such as Branin **(Supplementary Figure 1a)**, the performance gap between methods was small, as all approaches could locate near-optimal solutions efficiently. However, for more rugged and multimodal functions such as Ackley5 and Hartmann6 **(Supplementary Figure 1b and c)**, BO showed a clear advantage. In these cases, the probabilistic surrogate model and acquisition-driven search enabled BO, in both exploitative and explorative modes, to systematically identify high-performing regions. In contrast, LHS and Random Search frequently became trapped in local minima and failed to reach the global optimum within the same evaluation budget.

The bioprocess simulation group **(Supplementary Figure 1d–f)** includes surrogate functions such as perfusion_density_surrogate, titer_e, and viability_surrogate, which were constructed to mimic key features of real bioreactor systems. These functions incorporate process-inspired characteristics such as saturation effects, multiple performance basins, and experimental noise, capturing the nonlinear and uncertain behavior typical of upstream bioprocesses. Among these surrogates, the **perfusion_density_surrogate** in Figure 2d represents a pseudo– steady-state relationship between cell density and perfusion rate, incorporating diminishing returns and mild inhibitory behavior at high perfusion intensities. Bayesian Optimization (BO) rapidly identified high-performance regions within the first few iterations and showed consistent improvement in later iteration, whereas LHS and Random Search exhibited slower improvement and substantially higher variance across runs. The **titer_e** function (Supplementary Figure 2e) introduces a more rugged and multimodal landscape by combining nonlinear coupling between culture duration and productivity, and distance-dependent noise. This surrogate reflects the multi-factor interactions and stochastic variability commonly encountered in fed-batch and intensified processes. BO consistently outperformed the baseline methods, demonstrating both faster convergence and reduced variability when navigating this more challenging optimization surface. Finally, the **viability_surrogate** (Supplementary Figure 2f) captures viability decay dynamics driven by nutrient limitation and accumulation of inhibitory by-products, implemented through smooth saturation terms and a shallow interaction ridge. BO consistently achieved faster convergence, lower variance, and more reliable identification of high-performing regions than LHS and Random Search, despite increased noise and landscape complexity.

**Figure 2.**
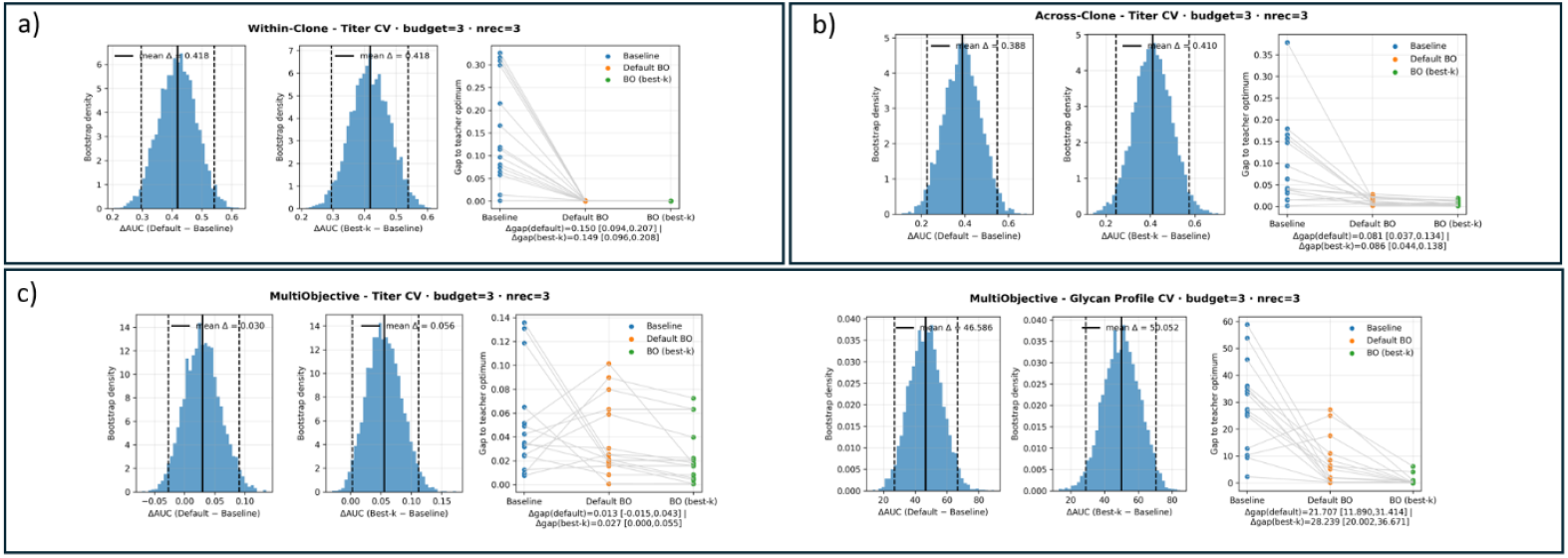
Offline-teacher evaluation across 3 scenarios. **a) Within-clone prediction. b) Clone/Scale Aware prediction. c) Multi-Objective Prediction**. Each row contains three panels per scenario. **Left:** bootstrap distribution of ΔAUC between the **default BO setting** (k = 1, nrec = 3) and the Random Search baseline; **Middle:** bootstrap distribution of ΔAUC between the **best-k BO setting** (selected per training seed) and the RS baseline. For **left** and **middle** panels, the X-axis represents ΔAUC and the Y-axis represents the sampling density. The sampling density reflects how frequently a given ΔAUC value appears across the 10,000 mean estimates derived from all fold–seed combinations, providing a visual summary of the overall performance shift between methods. The solid line indicates the mean and the dashed lines mark the 95% confidence interval (CI), and positive ΔAUC values indicate improved performance relative to the baseline. **Right:** paired scatter showing the gap to the teacher optimum for the RS baseline, the default BO, and the best-k BO for each evaluation run. The numbers printed underneath this panel report **Δgap**, defined as the average reduction in the gap to the teacher optimum relative to the baseline (Δgap = gap_baseline − gap_method), along with its 95% bootstrap CI. the best values achieved by the different methods within each fold–seed combination are connected by gray lines to facilitate direct visual comparison.

Overall, BO’s superior performance arises from its probabilistic modeling framework. By learning a Gaussian surrogate, BO captures both the expected response and the associated uncertainty across the design space. The Upper Confidence Bound (UCB) acquisition function leverages this uncertainty to adaptively balance exploration and exploitation, prioritizing experiments that are either highly promising or highly informative. Unlike space-filling or random approaches, BO learns from prior evaluations and focuses sampling on where it matters most. This enables rapid convergence with fewer evaluations and provides robustness to noise and nonlinearity, properties that are essential for optimization in biological and bioprocess systems.

### 3.2 Cross Validation on historical training sets

After benchmarking on analytic test functions, we evaluated the BO framework using historical CHO cell bioprocess data. The full dataset is summarized in **Table 2** and consists of 58 fed-batch runs from two production clones. Clone A produces a monoclonal antibody, while Clone B produces a bispecific antibody, representing both clone-to-clone and product-modality variability. The goal of this evaluation was to assess how well the optimizer learns from real bioprocess data and how its performance changes when moving from a controlled, clone-specific setting to a more realistic, heterogeneous training environment, while also establishing a robust evaluation framework for fast model testing and calibration without requiring new experimental data.

To this end, we constructed multiple training and validation scenarios using different subsets of the historical data. We compared within-clone and across-clone training strategies under three scenarios: (1) Within-clone, single-objective optimization, where the model was trained and evaluated exclusively on Clone A data. (2) Across-clone, single-objective optimization, where training data from Clone A and Clone B were combined to optimize Clone A performance. (3) Multi-objective optimization, where the model simultaneously optimized multiple objectives (e.g., titer and glycan profile) for Clone A using within-clone data from Clone A.

All scenarios used the grouped recommendation strategy, in which multiple candidate conditions are proposed per iteration by sampling across a range of length scales. This design supports parallel experimentation and reflects practical bioprocess workflows, where multiple conditions can be tested simultaneously rather than sequentially.

Model performance was evaluated using a five-fold teacher–student framework (Methods 2.2). In each evaluation, a teacher model defined the reference response surface and its optimum, while the optimizer (student) sequentially proposed experimental conditions under a fixed budget.

Performance was quantified using two complementary metrics (Figure 3):

- **ΔAUC**, defined as the difference in area under the convergence curve relative to the RS baseline. Higher ΔAUC indicates faster convergence toward the optimum.
- **Gap to the optimum**, defined as the difference between the best condition identified by the optimizer and the teacher-defined optimum. Smaller gaps indicate better final solutions.

**Figure 3.**
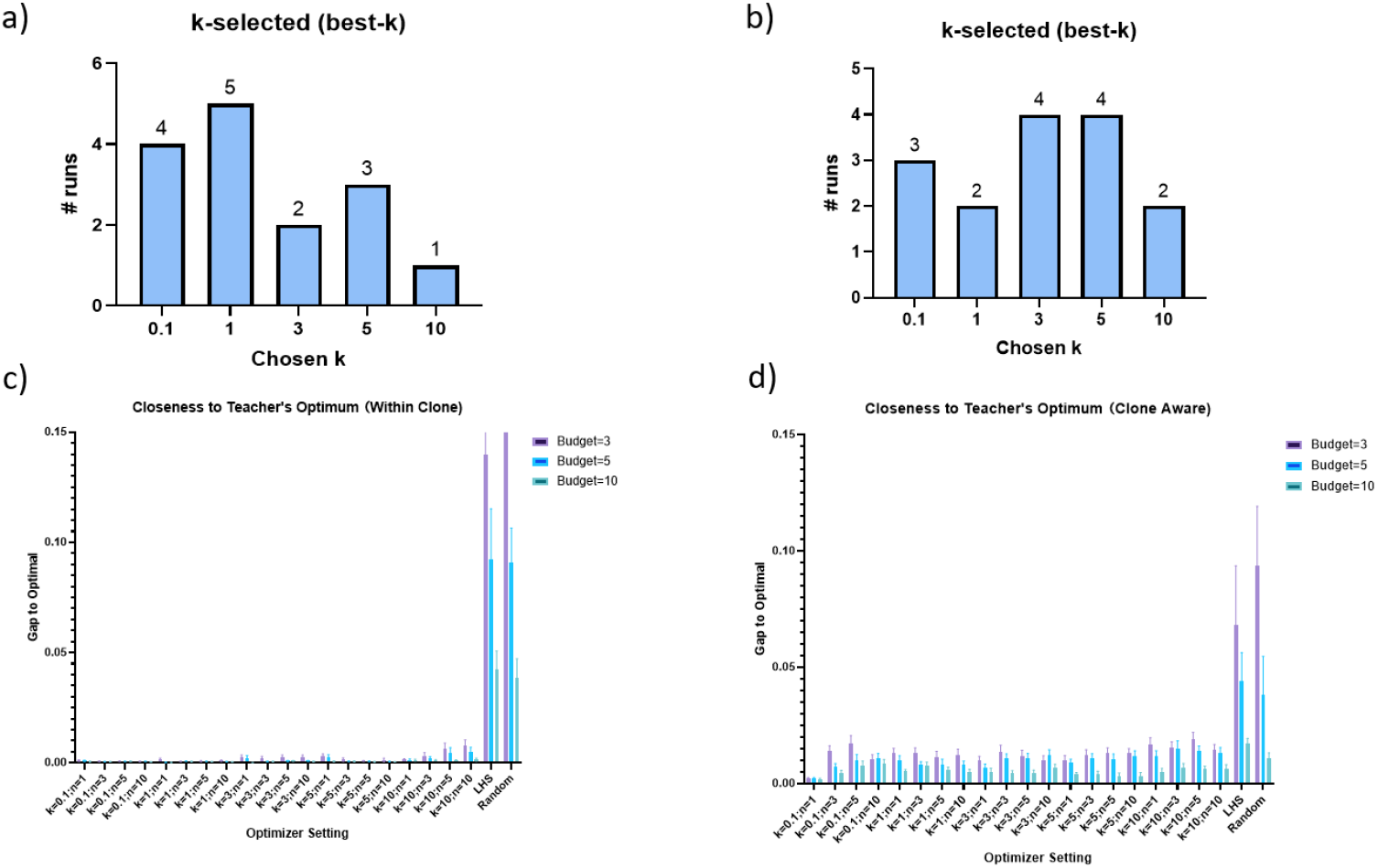
Comparison of Bayesian Optimization performance under within-clone and clone-aware training. (a–b) Distribution of the best-performing exploration–exploitation parameter (*κ*) across different random seeds and experimental budgets for (a) within-clone and (b) clone-aware models. Each bar represents the number of independent runs in which a given *κ*value yielded the best result. the y-axis indicates the number of fold-seed combinations selected at a given *κ* as the best performer, which produced the highest AUC within that run. (c–d) Corresponding optimization performance, shown as the mean gap to the teacher’s optimum under different budgets (1,3,5,10) for (c) within-clone and (d) clone-aware training. The Bayesian Optimizer achieved consistently low errors across all settings, with the within-clone model demonstrating slightly better accuracy and stability than the clone-aware configuration.

Across all scenarios, BO produced ΔAUC distributions centered above zero and consistently smaller gaps to the optimum than Random Search. Table 3 summarizes results across all evaluation settings, reporting the mean and standard deviation of the gap to the optimum for Random Search, default BO, and the best-performing BO configuration, defined as the *κ* value that yielded the smallest average gap to the optimum. In every case, BO outperformed Random Search, with the best-*κ* configuration achieving the closest match to the teacher model. In the multi-objective setting, BO improved both titer and glycan objectives, demonstrating its ability to balance competing goals.

**Table 3.**
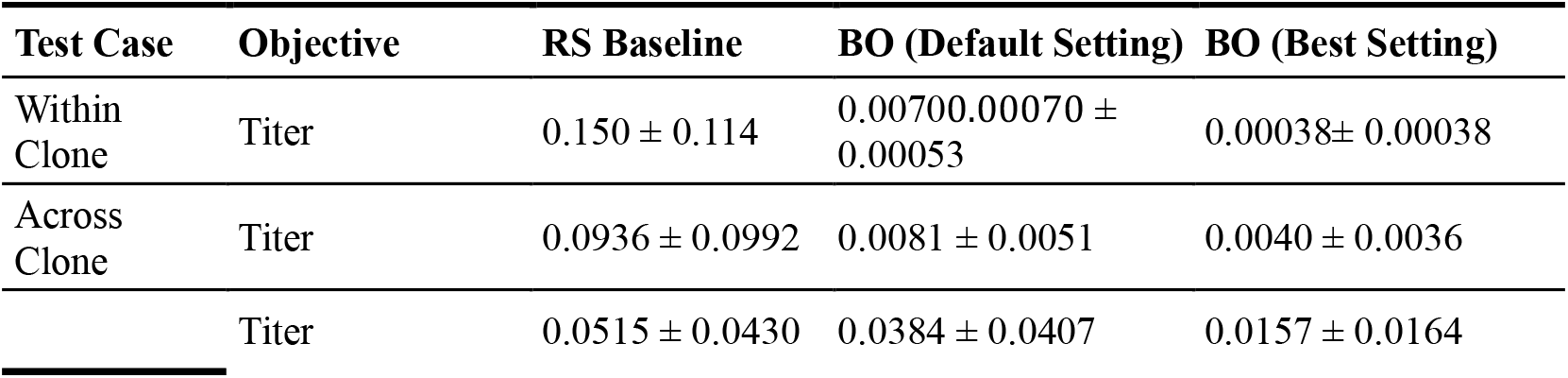
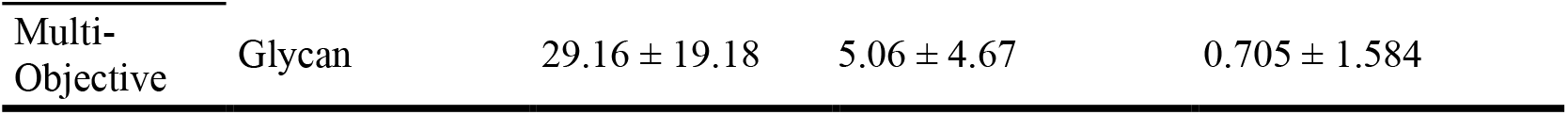
Comparison of model performance across three test scenarios: within-clone prediction, across-clone prediction, and multi-objective prediction. For each objective, the table reports the mean ± standard deviation of the gap to the teacher optimum achieved by the Random Search (RS) baseline, the default BO setting, and the best-performing BO setting. Smaller values indicate predictions closer to the teacher optimum.

| Test Case | Objective | RS Baseline | BO (Default Setting) | BO (Best Setting) |
| --- | --- | --- | --- | --- |
| Within Clone | Titer | 0.150 ± 0.114 | 0.00700.00070 ± 0.00053 | 0.00038± 0.00038 |
| Across Clone | Titer | 0.0936 ± 0.0992 | 0.0081 ± 0.0051 | 0.0040 ± 0.0036 |
| | Titer | 0.0515 ± 0.0430 | 0.0384 ± 0.0407 | 0.0157 ± 0.0164 |

#### 3.2.1 Within-clone evaluation

The within-clone evaluation measures optimizer performance in the most controlled setting, where both training and prediction use data from a single clone. This isolates the optimizer’s ability to exploit a well-defined, homogeneous response surface without confounding biological variability. We trained the model on Clone A, which had the largest historical dataset, and predicted optimal process parameters for the same clone. As shown in **Figure 2a**, the left and middle panels compare the ΔAUC distributions of the default BO configuration and the best-performing BO configuration against the LHS baseline. Both BO settings yield positive ΔAUC values with narrow distributions, indicating faster convergence than Random Search and high reproducibility across the five folds of the evaluation.

The right panel (gap-to-optimum) shows that both BO configurations consistently achieved smaller final gaps than Random Search across all 15 evaluations (five folds × three seeds). Quantitatively, BO reduced the average gap to the optimum by nearly two orders of magnitude relative to Random Search. These results demonstrate that under data-rich, clone-specific conditions, BO efficiently exploits learned structure to identify near-optimal process parameters.

#### 3.2.2 Across-clone evaluation

The across-clone evaluation assesses whether the optimizer can explore unknown solution space and generalize when trained on heterogeneous datasets spanning both distinct clones and different product modalities. In this clone- and molecule-aware setting, historical data from Clone A and Clone B in 250 mL were combined using hot encoding of categorical variables, and the model was tasked with optimizing Clone A titer. Clone A produces a monoclonal antibody, whereas Clone B produces a bispecific antibody, introducing variability not only in clonal background but also in product structure and processing demands. Providing clone identity and scale as categorical variables enables the optimizer to condition its response surface based on biological context, allowing it to learn similarities and sensitivities in process–performance trends across related but non-identical systems. This setup reflects realistic industrial scenarios where optimization knowledge must transfer across related but biologically non-identical processes.

As shown in **Figure 2b**, both the default and best BO configurations again achieved positive ΔAUC values relative to Random Search (mean ΔAUC = 0.388 and 0.41, respectively), indicating again faster convergence across most folds and seeds. The gap-to-optimum results show that BO reduced the final error in all but two cases. Although improvements were smaller than in the within-clone setting, this reduction is expected given the added uncertainty from inter-clone and molecule variability. Importantly, BO still consistently outperformed Random Search and identified near-optimal solutions, demonstrating robustness and effective knowledge transfer across heterogeneous bioprocess data.

#### 3.2.3 Multi-objective evaluation

We next extended the evaluation to a multi-objective setting, where the optimizer simultaneously targeted productivity (titer) and product quality (glycan profile). Training and prediction were performed within-clone on Clone A to isolate the effect of multi-objective trade-offs. This scenario reflects realistic bioprocess optimization challenges, in which improving one attribute often compromises another.

Results are shown in **Figure 2c**, with titer on the left and glycan profile on the right. For titer, BO achieved only modest improvements over Random Search, with mean ΔAUC values of 0.034 (default BO) and 0.056 (best-*κ* BO). These gains are smaller than in the single-objective cases, indicating limited remaining headroom for titer improvement under multi-objective constraints. In contrast, glycan optimization showed a strong and consistent improvement. The ΔAUC distributions are markedly right-shifted, with mean values of 46.586 and 50.052 for the default and best-*κ* BO configurations, respectively, corresponding to large reductions in glycan MSE relative to Random Search. Improvements were consistent across all fold–seed combinations.

Together, these results show that in a multi-objective setting, BO prioritizes improvement where flexibility exists (glycan profile) while maintaining near-optimal performance in more constrained objectives (titer), which is the desired behavior when objectives compete.

#### 3.2.4 Trends across experimental budgets and exploration strategies

To examine how exploration–exploitation balance and experimental budget affect performance, we evaluated BO across *κ* values (0.1, 1, 3, 5, 10) and batch sizes nrec (1, 3, 5, 10), spanning 12 experimental budgets.

**Figures 3a and 3b** show the *κ* values most frequently selected as optimal for within-clone and across-clone training, respectively, with nrec fixed at 3. Within-clone optimization favored smaller *κ* values (0.1–1), indicating that exploitative strategies are sufficient in stable, homogeneous landscapes. In contrast, across-clone optimization favored larger *κ* values (3– 10), reflecting the need for increased exploration under higher variability of mixed clone input.

**Figures 3c and 3d** summarize the gap between predicted and teacher-optimal titer at recommended conditions across budgets for within-clone and across-clone settings. In both cases, BO achieved consistently lower gaps to the optimum than LHS and random baselines across all *κ* values and budgets. The across-clone setting exhibited slightly higher gaps and variability, consistent with its greater complexity within the heterogeneous, across-clone dataset and aligns with the need of larger *κ* values shown in **Figure 3b**, but BO remained robust and convergent. Furthermore, increasing the experimental budget improved performance for all methods, however BO maintained a clear advantage at every budget level.

### 3.3 Forward experimental validation

Following in-silico benchmarking and retrospective evaluation on historical datasets, we next performed forward experimental validation to test whether the BO framework could prospectively improve bioprocess performance in real bioreactor experiments. These experiments directly assess whether the convergence and robustness observed in earlier sections translate into actionable process improvements.

The optimization variables included pH, dissolved oxygen (DO), temperature, and feed rate. Experimental outcomes were evaluated according to the relevant optimization objectives (titer, glycan profile, and charge variants), as summarized in **Table 1**. The normalized BO-selected operating conditions for each validation scenario are listed in **Table 4**, and all 21 reactor conditions tested across the study are provided in **Supplementary Table 2** for the entire explored space. A schematic overview of the experimental workflow is shown in **Figure 4a**.

**Table 4.** Summary of operating conditions and observed performance for the five Bayesian optimization recommendation cases. The table reports factor changes in pH, dissolved oxygen (DO), temperature, and feeding rate for each run, along with the associated optimization objective (titer, glycan MSE, charge variant MSE, or a combined titer/glycan objective) and the corresponding experimental output.

| Case # | pH | DO | Temperature | Feeding Rate | Objective |
| --- | --- | --- | --- | --- | --- |
| 1 | 1.02 | 1 | 1.04 | 1.02 | Titer |
| 2 | 1.03 | 0.40 | 1.09 | 0.89 | Titer |
| 3 | 1.03 | 0.53 | 1.09 | 0.5 | Glycan MSE |
| 4 | 1.05 | 1.33 | 0.85 | 1.08 | Charge Variant MSE |
| 5 | 1.04 | 0.41 | 1.01 | 0.97 | Titer/Glycan |

**Figure 4.**
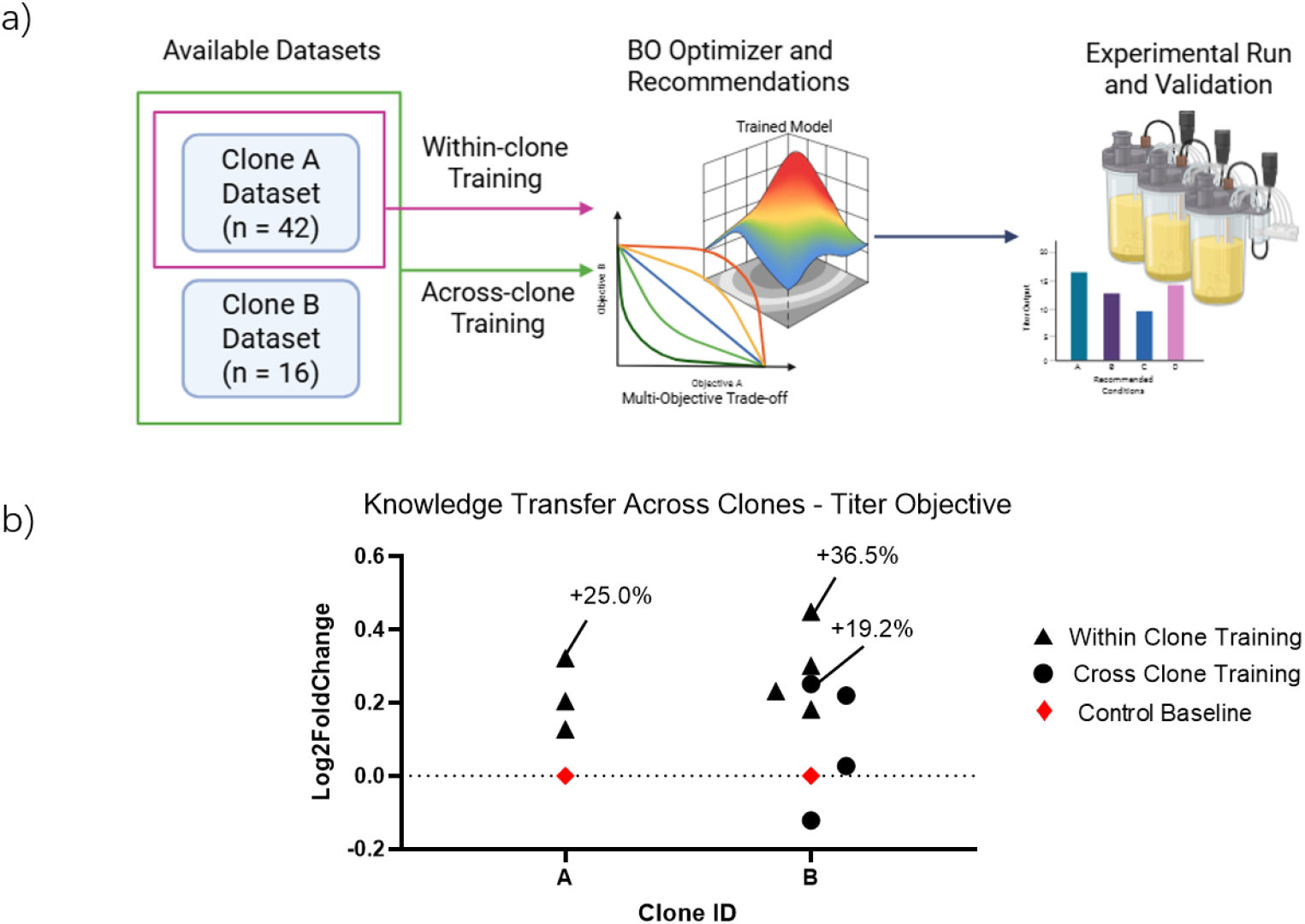
Within-clone and across-clone training strategies and their impact on titer optimization in CHO bioprocesses. **(a)** Schematic illustrating the within-clone and across-clone Bayesian optimization (BO) training strategies. Historical datasets from Clone A (n = 36) and Clone B (n = 16) were used to evaluate knowledge transfer across CHO clones. In the within-clone strategy, the BO model was trained and evaluated using data from a single clone, enabling learning of clone-specific process–performance relationships. In the across-clone strategy, data from both clones were pooled to train a unified BO model intended to capture shared biological and operational trends across clones. In both cases, trained models generated recommended operating conditions through single-objective or multi-objective optimization, which were subsequently implemented in bioreactor experiments for experimental validation. **(b)** Experimental validation of titer improvements under different BO training strategies. For clone 3F7, both within-clone (▲) and across-clone (●) BO models were evaluated under single-objective titer optimization, whereas for Clone A only the within-clone model was applied. Across all conditions, within-clone BO recommendations consistently achieved higher titers than across-clone recommendations, yielding up to **36.5%** and **25%** improvement over the control baseline (◆) under platform operating conditions.

Historical datasets (**Table 2**) from Clone A (n = 42) and Clone B (n = 16) were used to construct two training strategies: (i) a within-clone model, trained exclusively on Clone A data to capture clone-specific process–performance relationships, and (ii) an across-clone model, trained on the combined dataset to learn shared patterns across clone, molecule and scale.

Each trained BO model generated recommended process conditions under either single-objective or multi-objective formulations, which were then implemented experimentally.

#### 3.3.1 Forward validation of titer optimization

To test the generalizability and transferability of the optimization framework, we first evaluated within-clone and across-clone strategies for improving titer. In the within-clone setting, BO was trained and tested using data from the same clone, whereas in the across-clone setting, BO was trained on combined clone data and applied to a specific Clone. As shown in **Figure 4b**, the within-clone strategy was applied independently to Clone A and Clone B, reflecting the different sizes of their available historical datasets. For Clone A, all three within-clone test cases achieved at least a 10% increase in titer relative to the LHS baseline, with a maximum improvement of 25%. For Clone B, three of four within-clone test cases improved titer, with a maximum gain of 37% (black triangles).

The across-clone strategy was evaluated by training the BO on the combined Clone A and B dataset (58 total conditions) and applying to Clone B. Under this configuration, three of four test cases improved titer relative to the baseline, with a maximum increase of 19% (circles). While these gains confirm that transferable information across clones can support optimization, performance was consistently lower than in the within-clone case. This outcome aligns with earlier retrospective results and reflects the fact that clone-specific models better capture nuanced biological and process-level variability, and in particular when the product modality differs.

#### 3.3.2 Forward optimization of product quality attributes

Unlike titer maximization, product quality attributes such as glycan profiles and charge variants have defined compositional targets corresponding to acceptable and human-like post-translational modifications. These objectives were therefore formulated as error-minimization problems relative to predefined target setpoints (see Methods).

Quality optimization was performed for Clone B, with recommendations generated independently for glycan profile and charge variants. Because titer was not included as an objective, one of the three recommended conditions yielded insufficient antibody production and was excluded, leaving two evaluable conditions per objective. As shown in **Figure 5a**, the best BO-recommended conditions reduced mean squared error by 96% for charge variants and 92% for glycan profile relative to the control baseline. One charge-variant condition showed a slight degradation relative to baseline, reflecting the exploratory component of BO. When predictive uncertainty is high, exploration can occasionally sample suboptimal regions.

**Figure 5.**
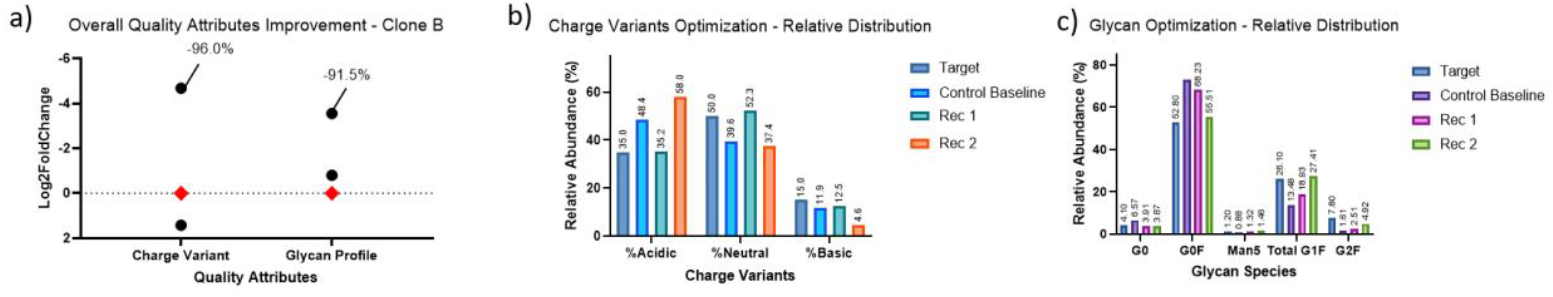
Optimization of product quality attributes on Clone B. (a) Overall quality improvement shown as log_2_-fold change relative to the control condition. Bayesian optimization reduced the charge-variant error by **96%** and glycan-profile error by **91.5%**, demonstrating strong convergence toward the target specifications. (b) **Distribution of acidic, neutral, and basic charge variants**. BO-recommended condition 1 shifted the profile toward the desired neutral-dominant composition closely and reduced off-spec acidic/basic species. (c) **Glycan-species distribution for the control baseline, target profile, and BO recommendations**. Both recommended conditions reduced high-mannose and non-fucosylated glycans while enriching target G0F/G1F species, resulting in a markedly closer match to the predefined target setpoint compared to the control baseline.

These outcomes are expected and informative, as they help refine the surrogate model and improve subsequent recommendations rather than indicating systematic failure.

To link these quantitative improvements to underlying biology, **Figures 5b** and **5c** break down species-level changes. For charge variants (**Figure 5b**), BO recommendations shifted the distribution toward the target composition by suppressing off-spec acidic and basic species and increasing the neutral fraction. For glycan optimization (**Figure 5c**), BO reduced high-mannose (Man5) and non-fucosylated glycans while enriching human-like structures such as G0F and G1F. The best recommendation minimized deviation across every monitored glycan class, consistent with the large reduction in glycan MSE. These results demonstrate that improvements arise from biologically meaningful shifts rather than compensatory effects.

#### 3.3.3 Multi-objective forward validation

Building on the single-objective results, we next evaluated the multi-objective BO module that simultaneously optimizes titer and glycan profile. This validation used the within-clone training strategy on Clone A, which provided the most extensive dataset.

As shown in **Figure 6**, the best BO recommendation achieved a 25% increase in titer while simultaneously reducing glycan profile MSE sevenfold (from 52.75 to 7.48). Importantly, all three multi-objective recommendations improved both objectives simultaneously, demonstrating that the BO framework effectively balances competing goals in a grouped experimental setting. These results directly mirror the retrospective findings and confirm that multi-objective gains are achievable in forward experiments.

**Figure 6.**
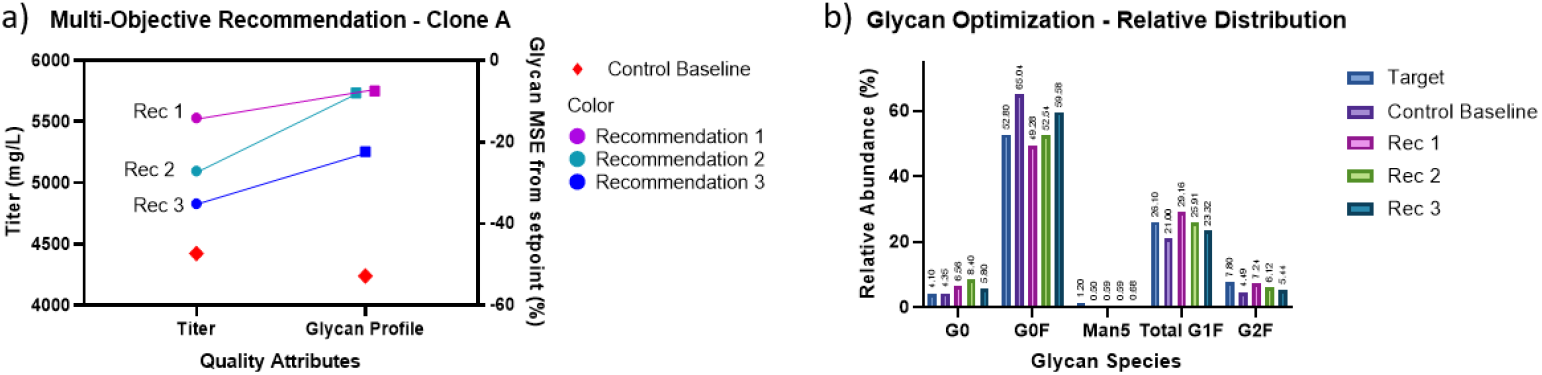
Multi-Objective Optimization (Titer/Glycan) on Clone A. a) Zoom-in on the clone A multi-objective recommendations (n = 3). The plot illustrates that titer improvement was accompanied by a simultaneous reduction in glycan profile mean-square error (MSE) relative to the setpoint, demonstrating that the proposed BO framework can enhance both productivity and product quality concurrently.b) Breakdown on the relative glycan abundance. All Recommendation showed shifts towards the setpoint.

#### 3.3.4 Biological interpretation of BO-recommended conditions

Analysis of the BO-recommended operating conditions revealed consistent and biologically interpretable trends linking process parameters to specific performance objectives. For titer optimization, BO recommendations for both Clone A and Clone B generally favored moderately elevated pH values (approximately 7.2–7.4) combined with intermediate temperatures (≈34.5–35.5 °C) and moderate feed rates, rather than aggressive feeding. These conditions are consistent with improved cellular productivity while limiting metabolic burden and byproduct accumulation. DO levels associated with higher titers tended to fall in a mid-to-upper range, suggesting sufficient oxygen availability to support oxidative metabolism without inducing excessive oxidative stress.

In contrast, optimization of charge variant and glycan quality attributes showed a clearer dependence on pH and temperature modulation. In particular, higher pH conditions were consistently associated with reduced charge-variant mean squared error, driven primarily by a shift toward the desired neutral species and suppression of acidic and basic variants. This trend aligns with known effects of extracellular pH on intracellular processing, including protein folding, deamidation, and charge heterogeneity^40,41^. Quality-optimized recommendations also favored slightly reduced temperatures and conservative feeding strategies, conditions that are known to stabilize post-translational modification pathways and reduce stress-induced heterogeneity^42^. Notably, Clone A and Clone B exhibited distinct response patterns, reflecting differences in both clone background and product modality. Clone A (monoclonal antibody) showed more pronounced titer gains under modest temperature reductions and balanced feeding, whereas Clone B (bispecific antibody) displayed greater sensitivity of quality attributes to pH adjustments, particularly for charge variants. These differences underscore the added complexity of molecule-specific processing demands and highlight why clone- and molecule-aware optimization is necessary for robust process development. Together, these results demonstrate that the BO framework identifies numerically optimal conditions that converge on mechanistically plausible operating regimes that reflect established relationships between pH, oxygen availability, feeding intensity, and CHO cell productivity and product quality.

## 4 Discussion

This study demonstrates a BO framework capable of optimizing CHO bioprocess performance across multiple objectives, clones, and scales. By integrating probabilistic modeling with an acquisition-driven exploration-exploitation strategy, the BO approach consistently outperformed conventional baseline process settings for industrial processes, even under tight experimental budgets. Building on our previous work in BO, we have shown that the same framework can go beyond basal media design to handle both continuous process variables (e.g., pH, dissolved oxygen, temperature, and feed rate) and categorical factors (e.g., clone identity and scale). This expansion enables the algorithm to capture biological and operational variability across different production systems, which is an essential step toward data-driven generalization in biomanufacturing^43^. In addition, the algorithm is designed to generate multiple recommendations per batch, enabling exploration of several promising regions of the design space simultaneously and aligning well with bioprocess workflows that rely on parallel experimentation.

The framework was validated in silico, on historical data, and followed by experimental verification of forward predictions. Across all these settings, the BO algorithm achieved faster convergence and smaller gaps to the pseudo-optimum, and higher overall process yields especially under limited experimental budgets. The comparison between training using data from a single clone (within-clone model) and training using data from multiple clones (across-clone model) highlights the difficulty of transfer learning across different clones and product modalities. In this study, Clone A produces a monoclonal antibody, whereas Clone B produces a bispecific antibody, introducing differences in folding, secretion, and quality-attribute sensitivity in addition to clonal variability^44^. Although both approaches improved titer relative to the control, the within-clone model consistently outperformed the across-clone training model in both historical dataset training and experimental validation. This drop in across-clone performance likely arises from the uneven historical dataset sizes. Clones with larger, more diverse data (Clone A) exhibited greater influence during model fitting, causing predictions to skew toward data-rich conditions and reducing generalizability to underrepresented clones. This can be addressed by adopting imbalance-aware modelling strategies that prevent data-rich clones from dominating the learning process. Approaches such as weighted loss functions or oversampling can help rebalance the training distribution so that underrepresented clones contribute more meaningfully to model fitting. In addition, incorporating clone-specific latent representations or hierarchical architectures that capture underlying biology, such as baseline phenotype and omics data, would allow the model to learn shared global trends while still preserving clone-level differences in productivity and process response. Together, these strategies can improve cross-clone generalizability and strengthen the reliability of transfer learning when applying a single model across diverse cell lines and molecules. In addition, the present study does not assess the ability of the model to transfer learned knowledge to a completely unseen clone. While the across-clone training results demonstrate partial knowledge sharing across different production systems, both clones used in this work were observed during training. Generalization to unseen clones remains an important open question. Addressing this will require larger and more diverse multi-clone datasets. Incorporating clone-aware representations, hierarchical modelling, or phenotype-informed priors may improve the model’s ability to extrapolate beyond previous observed cell lines.

A major strength of our new framework lies in its ability to optimize multiple, biologically coupled objectives simultaneously. CHO cell bioprocess optimization traditionally faces a fundamental trade-off where increasing titer can compromise product quality attributes, whereas targeting glycan quality can push cultures into conditions that produce insufficient material for meaningful assessment^45,46^. We observed this directly when optimizing solely for glycan profile, the resulting condition dropped out of downstream evaluation due to extremely low titer, illustrating the risk of pursuing quality objectives in isolation. Conversely, when optimizing exclusively for titer, we achieved a 36% increase in product titer but observed substantial deviations in glycan patterns, confirming that maximizing productivity alone does not ensure acceptable quality attributes. Our multi-objective formulation avoided these pitfalls by jointly modelling and balancing both objectives. By allowing the algorithm to navigate the nonlinear trade-off surface between productivity and quality, we achieved a 25% increase in overall titer and reduced glycan mean-square error by more than sevenfold under multi-objective targeting. This demonstrates that the framework not only prevents failure modes inherent to single-objective optimization but also identifies Pareto-efficient conditions that improve both yield and quality simultaneously. This capability enables more reliable, biologically meaningful optimization in complex CHO cell culture landscapes where competing objectives cannot be decoupled.

Although in-silico benchmarking studies do not fully replicate how the proposed BO framework is deployed in real multi-batch, multi-objective bioprocess optimization, these benchmarks provide a controlled and reproducible environment for validating the core algorithmic components. This framework includes convergence behavior, exploration-exploitation trade-offs, and robustness while using limited evaluation budgets. Establishing strong performance across a diverse set of analytic and surrogate benchmarks therefore serves as a critical testing ground, supporting and validating both the current implementation and its extensibility to more complex, real-world optimization scenarios.

Overall, this study establishes a multi-objective BO framework that captures both quantitative performance metrics and qualitative product attributes. The gains in titer and the reductions in glycan-profile error across multiple experimental conditions highlight the value of BO as a practical and data-efficient tool for CHO cell bioprocess development. This approach aligns with the growing interests toward autonomous, AI-driven experimental design in life sciences. When coupled with automated high-throughput cell culture systems, this multi-objective BO framework can accelerate process discovery cycles, enabling adaptation to emerging production challenges, new modalities and production hosts. Such integration of machine learning with bioprocess knowledge and automated systems creates a scalable, data-efficient workflow, and coupling it with uncertainty-based algorithms provides a powerful optimization engine for advancing biomanufacturing workflows.

## 5 Conclusion

This work demonstrates how AI-guided bioprocess development through BO can help improve biomanufacturing pipelines by providing a principled, uncertainty-aware framework for navigating the complexity of CHO cell manufacturing. As biologics continue to diversify, spanning multispecific antibodies, complex glycoengineered modalities, and cell-based therapies, traditional process-development strategies are becoming increasingly resource-intensive and difficult to scale. Our results highlight how BO can reduce this burden by making each experiment more informative, enabling rapid hypothesis generation, and guiding exploration toward process conditions that balance productivity and product quality attributes.

In addition to early-stage development, this approach has direct relevance for Process Characterization Studies (PCS), where efficient identification of critical process parameters and operating ranges is essential. By prioritizing experiments that maximally reduce uncertainty, BO can help reduce the number of runs required to establish process understanding and robustness, supporting more focused and data-efficient PCS execution. When deployed alongside emerging automation technologies such as robotic liquid handlers, integrated ambr® platforms, and real-time PAT sensors, the framework presented here can support closed-loop experimentation where models and experiments iteratively refine one another. Such an optimization framework would allow organizations to accelerate PCS, shorten tech-transfer timelines, and respond more quickly to shifts in molecular pipelines or manufacturing demands^7,47^. In this way, AI-driven optimization serves as a computational tool for next-generation biomanufacturing that is adaptive, scalable, and capable of meeting global therapeutic needs with greater speed and efficiency.

## Acknowledgments

The authors would like to thank Lonza Biologics Plc for funding the project work and the Lonza SMEs for manuscript review.

## Author Contributions

**E.M**. conceptualized the study, designed and implemented the core algorithmic framework, performed computational analyses, interpreted the results, and drafted and revised the manuscript. **J.M**. Interpreted the results, and drafted and revised the manuscript. **S.D**. and **Z.L**. performed computational analyses, interpreted the results, and drafted and revised the manuscript. **S.R**., **D.D**. conceptualized the study, designed and coordinated the lab experimentation, performed process analyses, interpreted the results, drafted and revised the manuscript. **C.L**., **S.A**., **A.D**. performed 15 mL experimentation and data analysis. **AG.G**. Conceptualized the 15 mL experimental design and provided inputs on the manuscript and figures. **L.K.W.F, M.S**. provided analytical support for product characterization assays. **P.V**. provided inputs to study and result interpretations. **M.J.B**., supervised the study, interpreted the results, and drafted and revised the manuscript.

## Data Availability

Cell growth and metabolite secretion curves are provided in Supplementary Data

## Code Availability

The algorithm was developed under an industrial-academic contract and is subject to contractual restrictions. The code is hosted in a private repository and may be made available to qualified researchers for non-commercial purposes upon reasonable request and approval. Additional information required to evaluate or reproduce the analyses reported in this article is available from the lead contact upon request, subject to applicable confidentiality and contractual obligations.

