## Supplementary Document for "Multi-Objective Bayesian Optimization for Data-Efficient Bioprocess Development"

### Supplementary file

**Supplement Table 1. In-silico test cases.** Fifteen objectives spanning convex, classic multimodal, medium/high-dimensional, noisy inputs, and bioprocess-relevant surrogates (Double Langmuir, Haldane–Monod, Chemostat-SS, Titer-A/B/E). Columns list input dimensions, domains, and empirical equations. All problems are cast to minimization.

| # | Name | Dim. | Inputs & Domain | Formula | Description |
|---|---|---|---|---|---|
| 1 | Paraboloid | 2 | $(x, y) \in [-5, 5]^2$ | $f(x, y) = x^2 + y^2$ | Simple convex "bowl" with a single global optimum at (0,0)(0,0). |
| 2 | Branin | 2 | $x \in [-5, 10], y \in [0, 15]$ | $f(x, y) = (y - (5.1/(4\pi^2)) \cdot x^2 + (5/\pi)x - 6)^2 + 10(1 - 1/(8\pi)) \cdot \cos(x) + 10$ | Two-dimensional, three global minima, widely used in BO benchmarks. |
| 3 | Hartmann-6 | 6 | $x \in [0,1]^6$ | $f(x) = -\Sigma_{i=1}^{4} \alpha_i \cdot \exp(-\Sigma_{j=1}^{6} A_{ij} \cdot (x_j - P_{ij})^2)$ $\alpha = [1.0, 1.2, 3.0, 3.2]$ | Six-dimensional multimodal function with several local minima. |
| 4 | Ackley-5 | 5 | $x \in [-32.768, 32.768]^5$ | $f(x) = -20 \cdot \exp(-0.2 \cdot sqrt((1/5) \cdot \Sigma_i x_i^2)) - \exp((1/5) \cdot \Sigma_i \cos(2 \cdot \pi \cdot x_i)) + 20 + e$ | Many small local minima in a nearly flat outer region |
| 5 | Rastrigin-5 | 5 | $x \in [-5.12, 5.12]^5$ | $f(x) = 10 \cdot 5 + \sum_{i=1}^{5} [x_i^2 - 10 \cdot \cos(2 \cdot \pi \cdot x_i)]$ | Regular grid of local minima, highly periodic. |
| 6 | Noisy Branin | 2 + noise | same as Branin + noise | $f(x, y) = Branin(x, y) + \varepsilon, \quad \varepsilon \sim Normal(0, 0.1^2)$ | Branin with homoscedastic Gaussian noise on observations. |
| 7 | Six-Hump Camelback | 2 | $(x_1, x_2) \in [-3, 3] \times [-2, 2]$ | $f(x_1, x_2) = \left(4 - 2.1x_1^2 - \frac{x_1^4}{3}\right)x_1^2 + x_1x_2 + (-4 + 4x_2^2)x_2^2$ | Classic 2-D multimodal surface with six local minima; two globals at (±0.0898, ±0.7126, $f_{min} \approx -1.0316$ |
| 8 | Hartmann-3 | 3 | $x \in [0,1]^3$ | $f(x) = -\sum_{i=1}^{4} \alpha_i \exp(-\sum_{j=1}^{3} A_{ij}(x_j - P_{ij})^2); \alpha = [1.0,1.2,3.0,3.2]$ | 3-D variant with a single global basin; $f_{min} \approx -3.8628$ near (0.115,0.556,0.853) |
| 9 | Double Langmuir | 1 | $c \in [0,10]$ | $f(c) = \frac{q_{1,max}K_1c}{1 + K_1c} + \frac{q_{2,max}K_2c}{1 + K_2c}$ | Two-site adsorption isotherm; smooth, saturating capacity vs. concentration. |
| 10 | Haldane–Monod (substrate inhibition) | 1 | $S \in [0,50]$ | $f(S) = \mu_{max}\frac{S}{K_S + S + \frac{S^2}{K_I}}$ | Growth rate with inhibition at high SS; interior optimum at $S = \sqrt{K_SK_I}$ |
| 11 | Titer-A (Lon×Prod) | 2 | $x \in [0,10]\ (longevity), y \in [0,5]$ | $f(x, y) = (1 - \exp(-0.3x)) \times (0.5y)$ | A smooth, monotonic function modeling "titer = longevity × productivity." |
| 12 | Titer-B (Interaction) | 2 | same as Titer-A | $f(x, y) = (1 - \exp(-0.3x)) \times (0.5y) + 0.2 \cdot \sin(0.8xy)$ | Adds a sinusoidal interaction term to the basic titer function. |
| 13 | Titer-E (Noisy) | 2 + noise | $x \in [0,10]^2$ | $f(x) = -(\sum_{k=1}^{K} a_k \exp\left(-\frac{\lVert x - c_k \rVert^2}{2w_k^2}\right) + g(x) + \epsilon(x)$ | Multi-basin sinusoidal titer surrogate with heteroscedastic noise |
| 14 | Perfusion Surrogate | 2 + noise | $x \in [0,10]^2$ | $f(x) = -(B(x) + S(x) + I(x) - P(x) + \epsilon)$, $B(x)$: *broad plateau*; $S(x)$: *local productivity peak*; $I(x)$: *Interaction term*; $P(x)$: *high density penalty* | Perfusion cell-density surrogate with plateau behavior, narrow high-performance pocket, and transport-limited penalty |

### Supplementary Figure 1

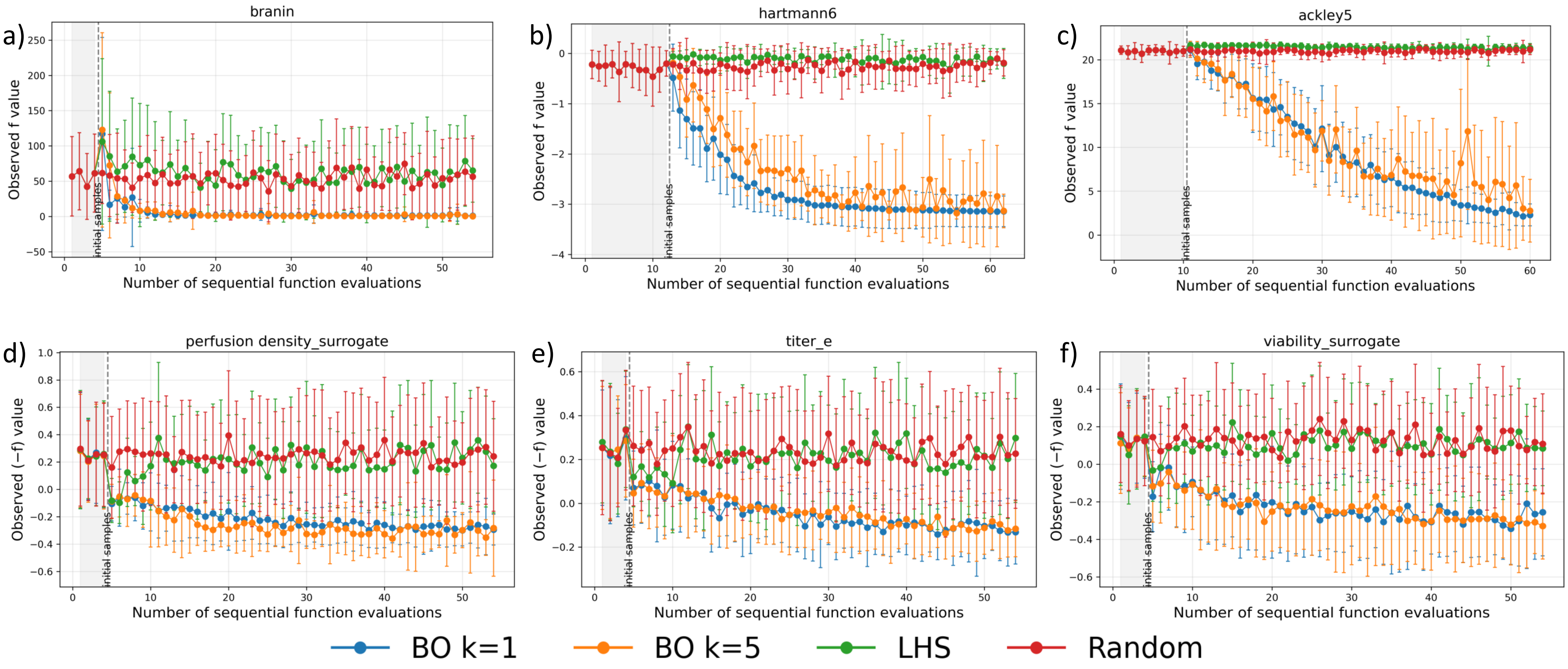


**Supplementary Figure 1. Performance over iteration on selected in-silico benchmarks.** Panels (a-f) plot the best-so-far objective versus number of function evaluations for BO (kappa=1), BO (kappa=5), Latin Hypercubic Sampling and Random Sampling on ackley5, hartmann6, rastrigin5, chemostat steady state simulation and two simulated titer cases. Shaded regions mark the initial samples; the vertical dashed line marks the start of adaptive search. Error bars show mean ± 1SD. Negative objective was plotted for maximization problem. Across the panel, BO consistently improves earlier and to lower final values than LHS/Random; BO with k=1 typically makes the fastest progress.

### Supplementary Table 2

| Running Clone | Training Method | Objective | pH | DO | Feed Rate | Temperature |
|---|---|---|---|---|---|---|
| Clone B | Across-Clone | Titer | 1.04 | 0.39 | 0.86 | 1.04 |
| | | | 1.05 | 1.05 | 0.96 | 1.09 |
| | | | 1.05 | 0.94 | 0.99 | 1.05 |
| | | | 1.02 | 1.33 | 0.94 | 1.05 |
| | | | 1.04 | 1.30 | 0.5 | 1.08 |
| | Within-Clone | Titer | 1.05 | 1.12 | 1.01 | 1.01 |
| | | | 1.03 | 0.40 | 0.89 | 1.09 |
| | | | 1.03 | 1.00 | 1.02 | 1.02 |
| | | | 1.02 | 1.00 | 1.02 | 1.04 |
| | | | 1.02 | 1.00 | 1.07 | 1.00 |
| | | Glycan Profile | 0.97 | 0.33 | 0.68 | 1.06 |
| | | | 1.04 | 0.53 | 0.5 | 1.09 |
| | | | 1.05 | 0.50 | 0.52 | 1.09 |
| | | Charge Variant | 1.03 | 1.33 | 1.49 | 0.85 |
| | | | 1.05 | 1.33 | 1.08 | 0.85 |
| | | | 1.05 | 1.18 | 1.5 | 0.91 |
| Clone A | Within-Clone | Titer & Glycan (Multi-Objective) | 1.04 | 0.41 | 0.97 | 1.01 |
| | | | 1.02 | 1.33 | 0.76 | 0.97 |
| | | | 1.05 | 0.37 | 0.94 | 1.01 |
| Clone A and B | Control Baseline | N/A | 1.00 | 1.00 | 1.00 | 1.00 |

Figure 1 : ambr15 DoE space

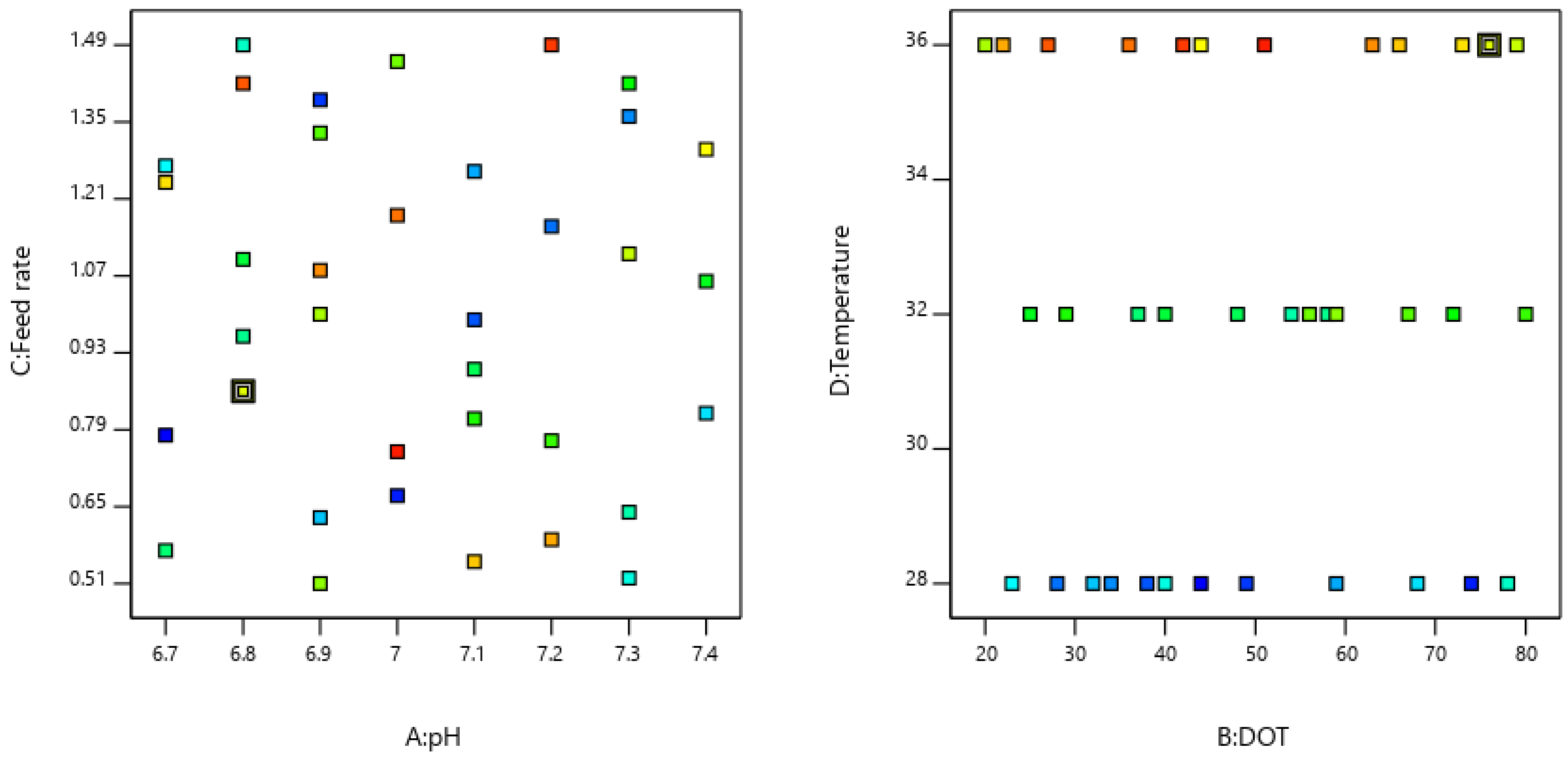

Figure 2 : Viable cell concentration profile for Clone A across different process parameter conditions in 15 mL vessels across different culture days

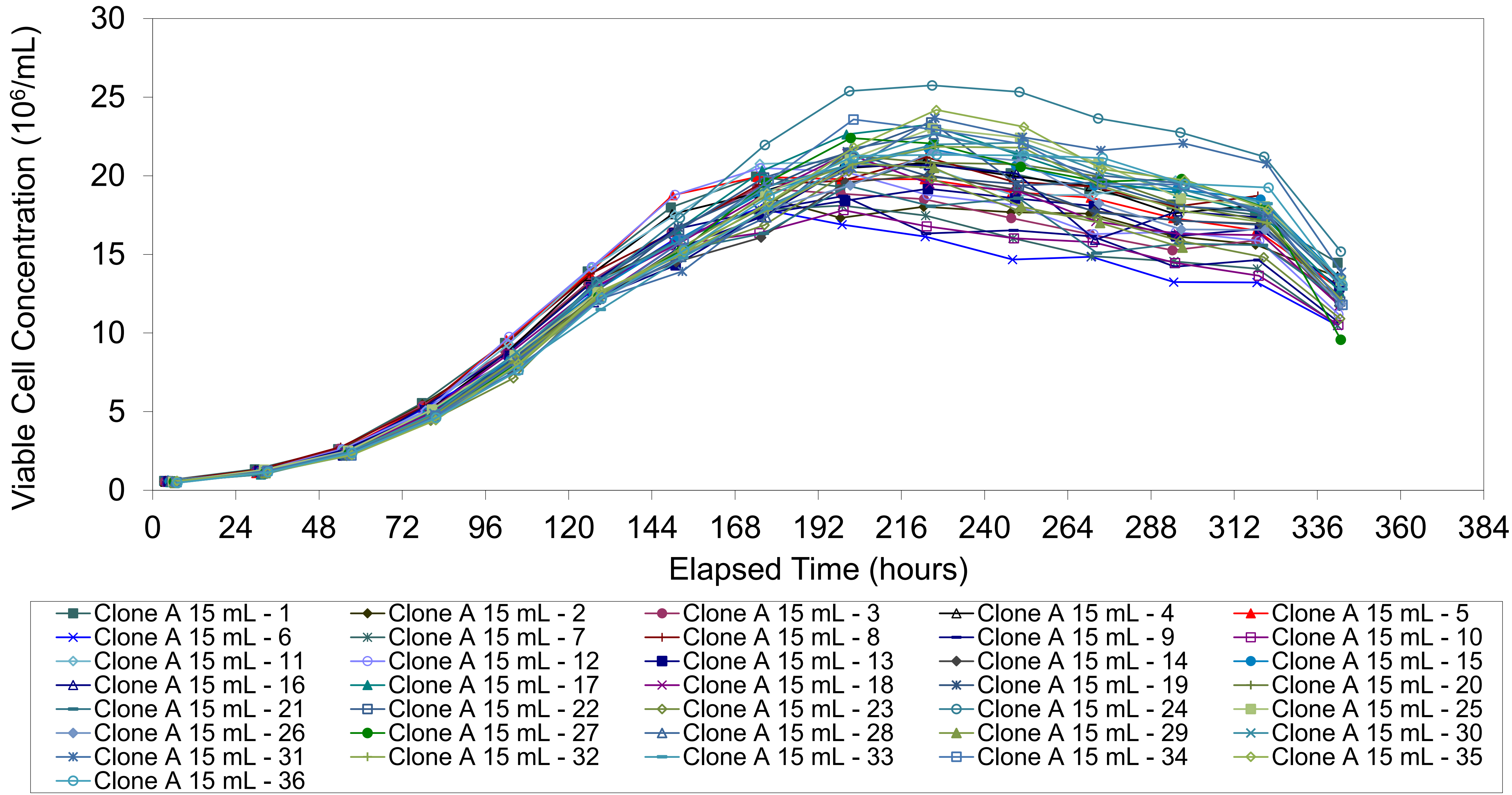

Figure 3 : Viability profile for Clone A across different process parameter conditions in 15 mL vessels across different culture days

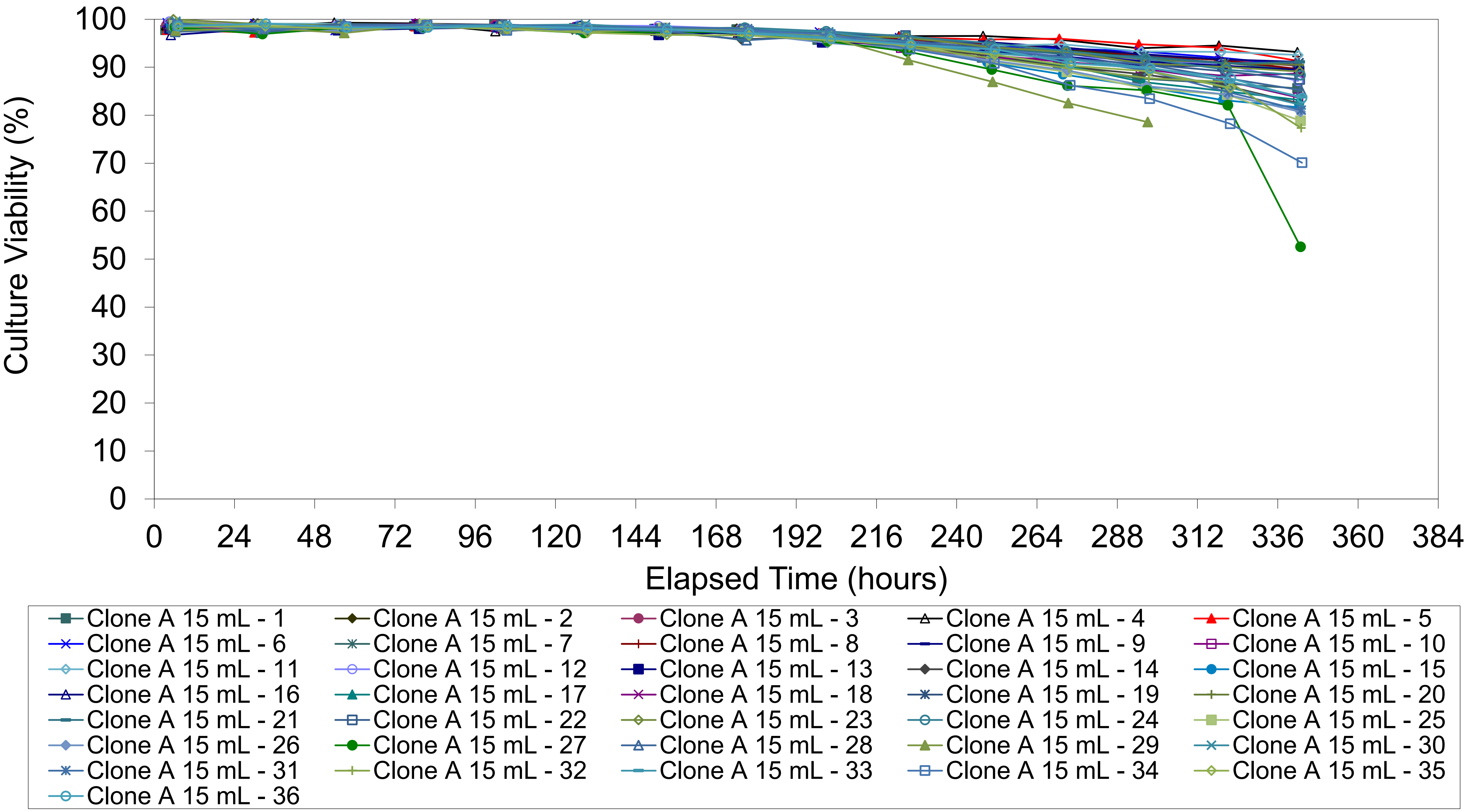

Figure 4 : Lactate concentration for Clone A across different process parameter conditions in 15 mL vessels across different culture days

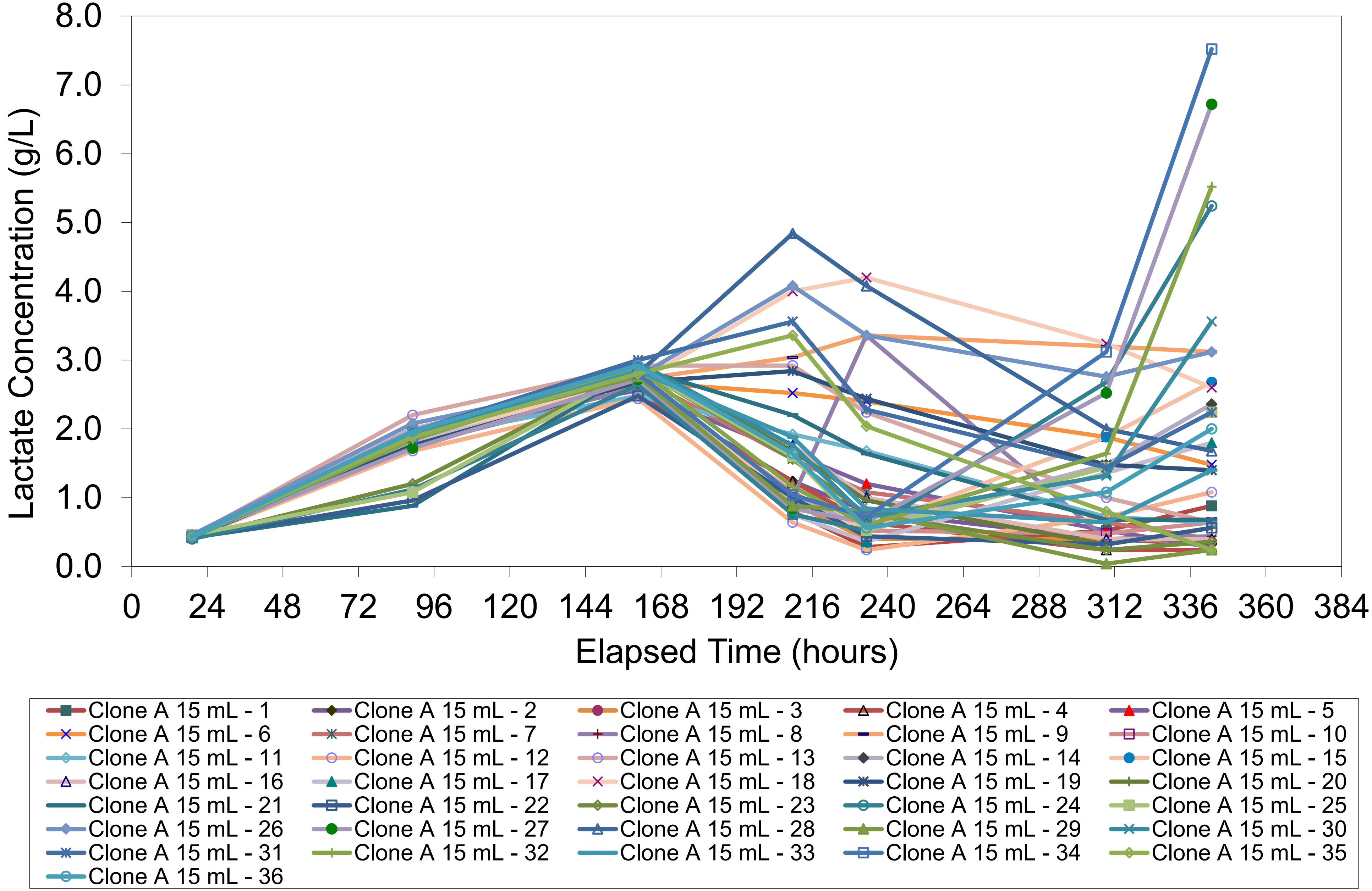

Figure 5 : Ammonium concentration for Clone A across different process parameter conditions in 15 mL vessels across different culture days

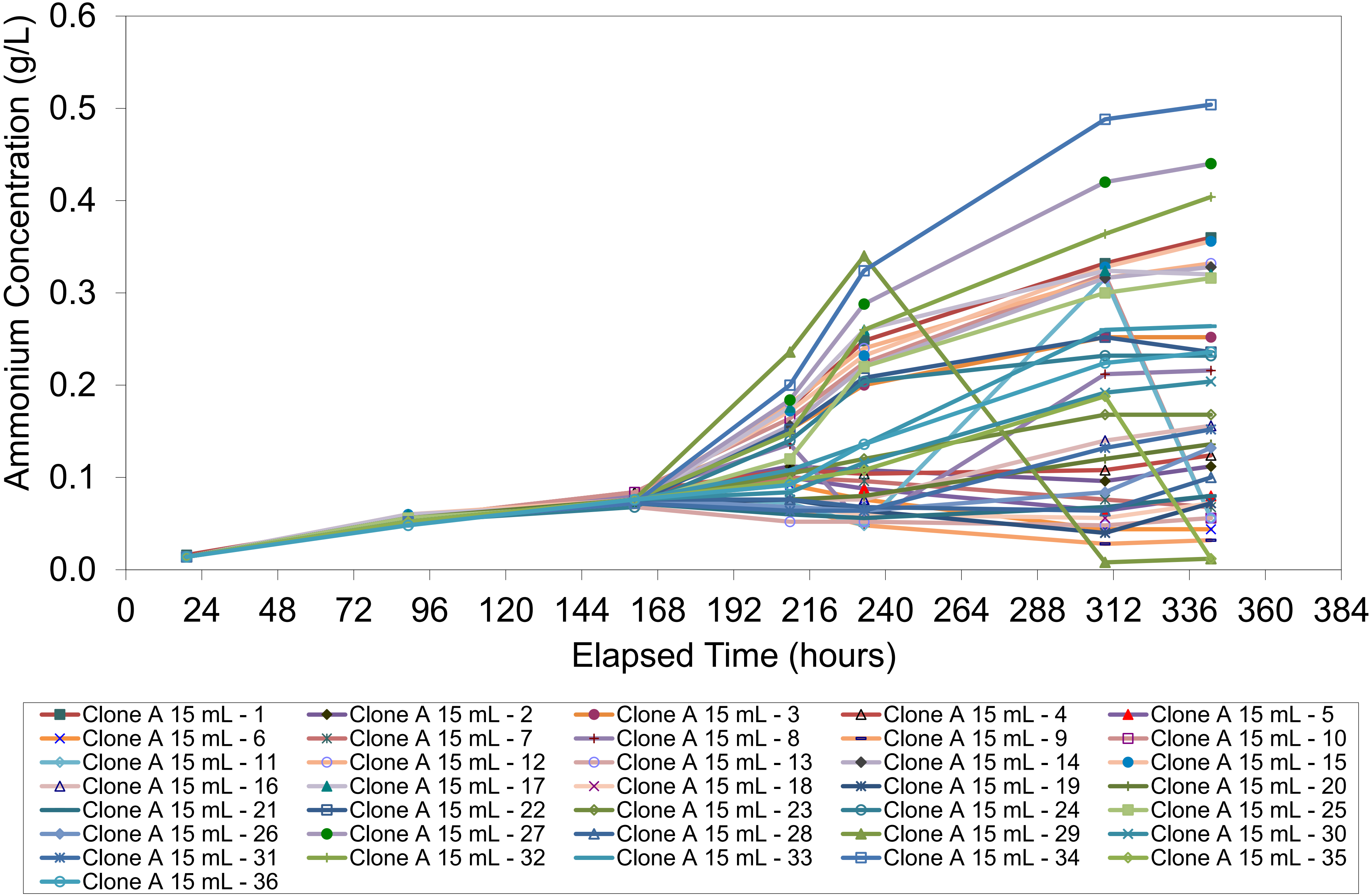

Figure 6a,b

Harvest titer , % agalactosylated antibody variants profile across clone type and scales

Titer

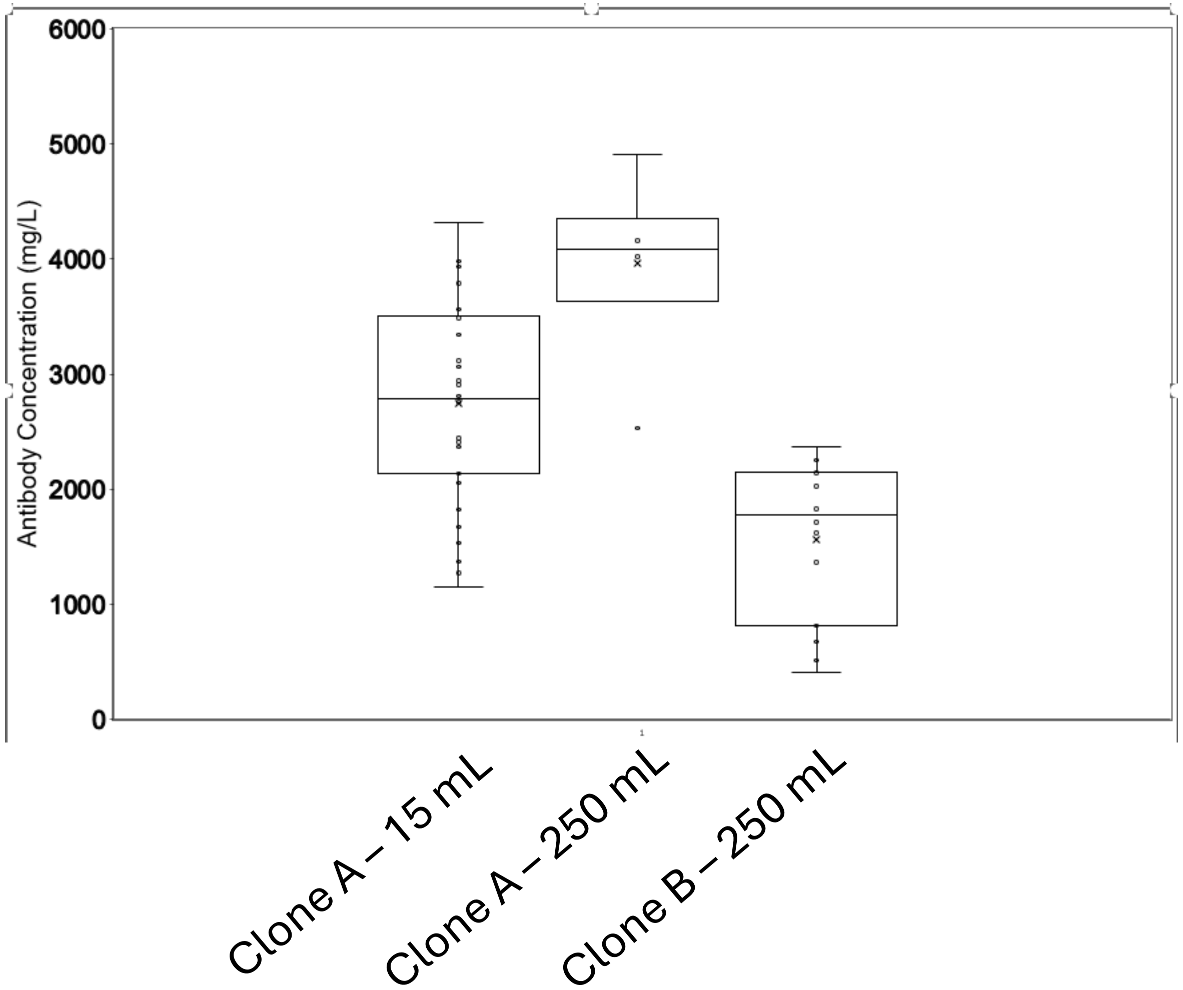


% Glycoform variants

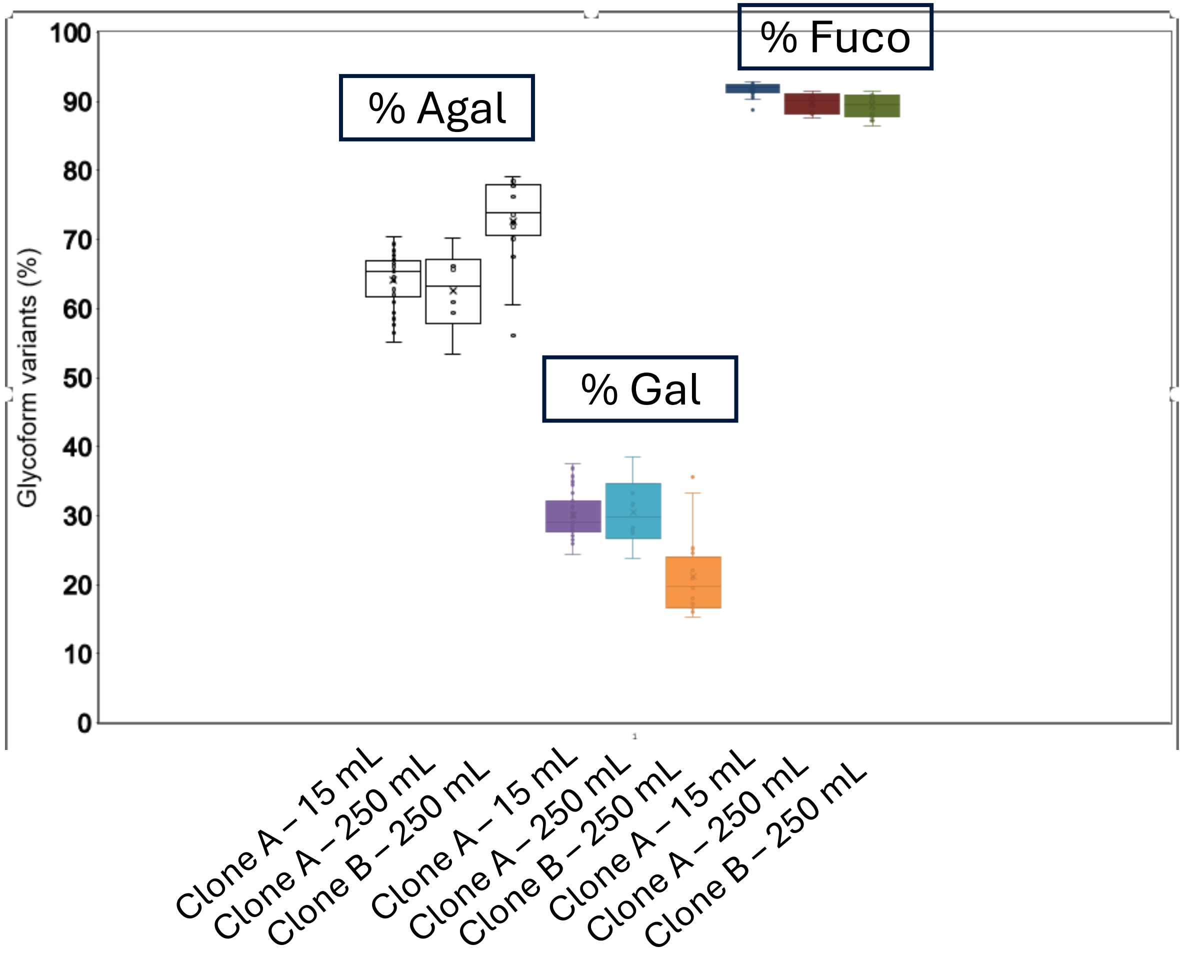




CONFIDENTIAL

Figure 6c

### % charged antibody variants profile across clone type and scales

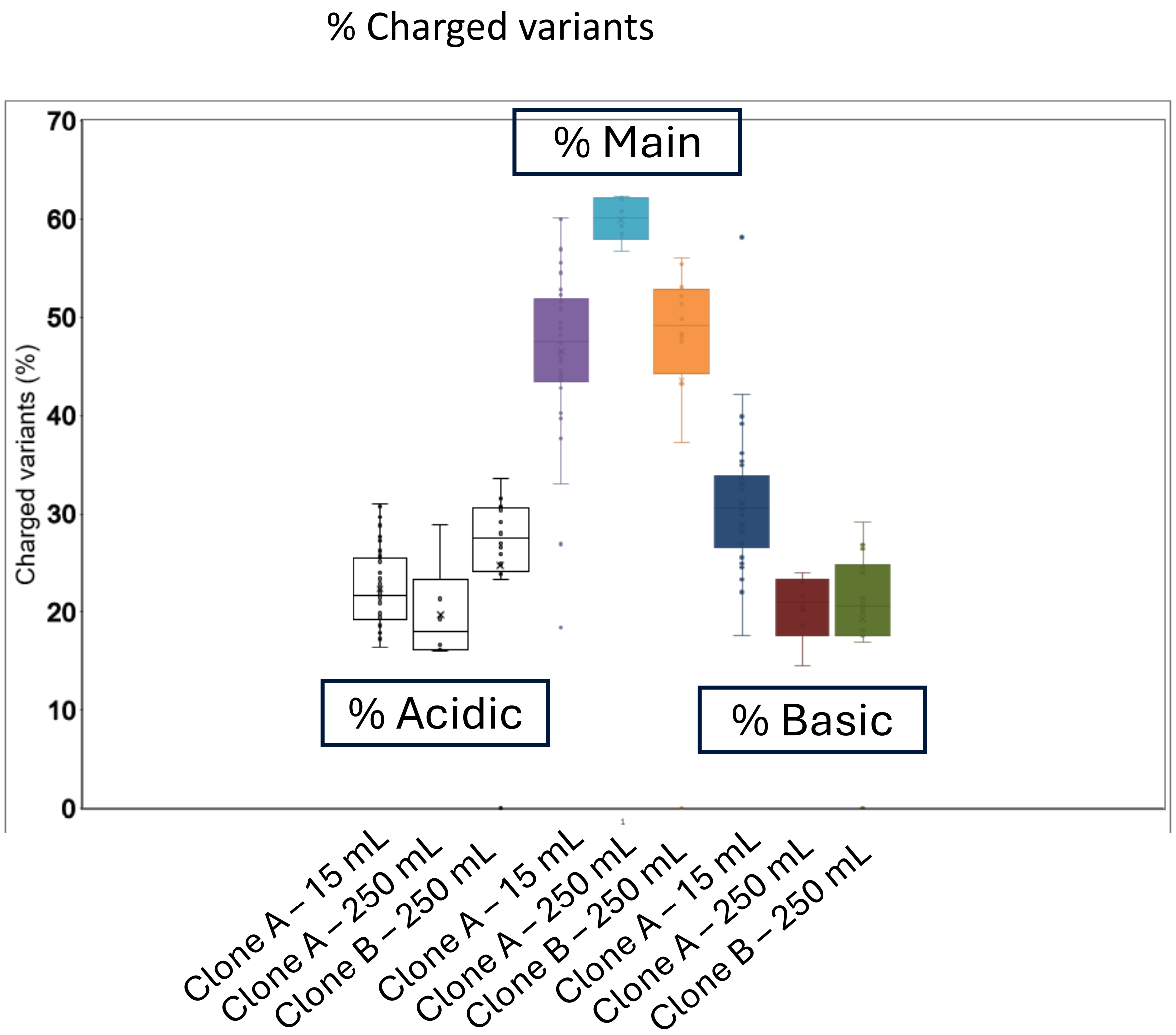




CONFIDENTIAL

Figure 3 : Viable cell concentration profile for Clone A & B across different process parameter conditions in 250 mL vessels across different culture days

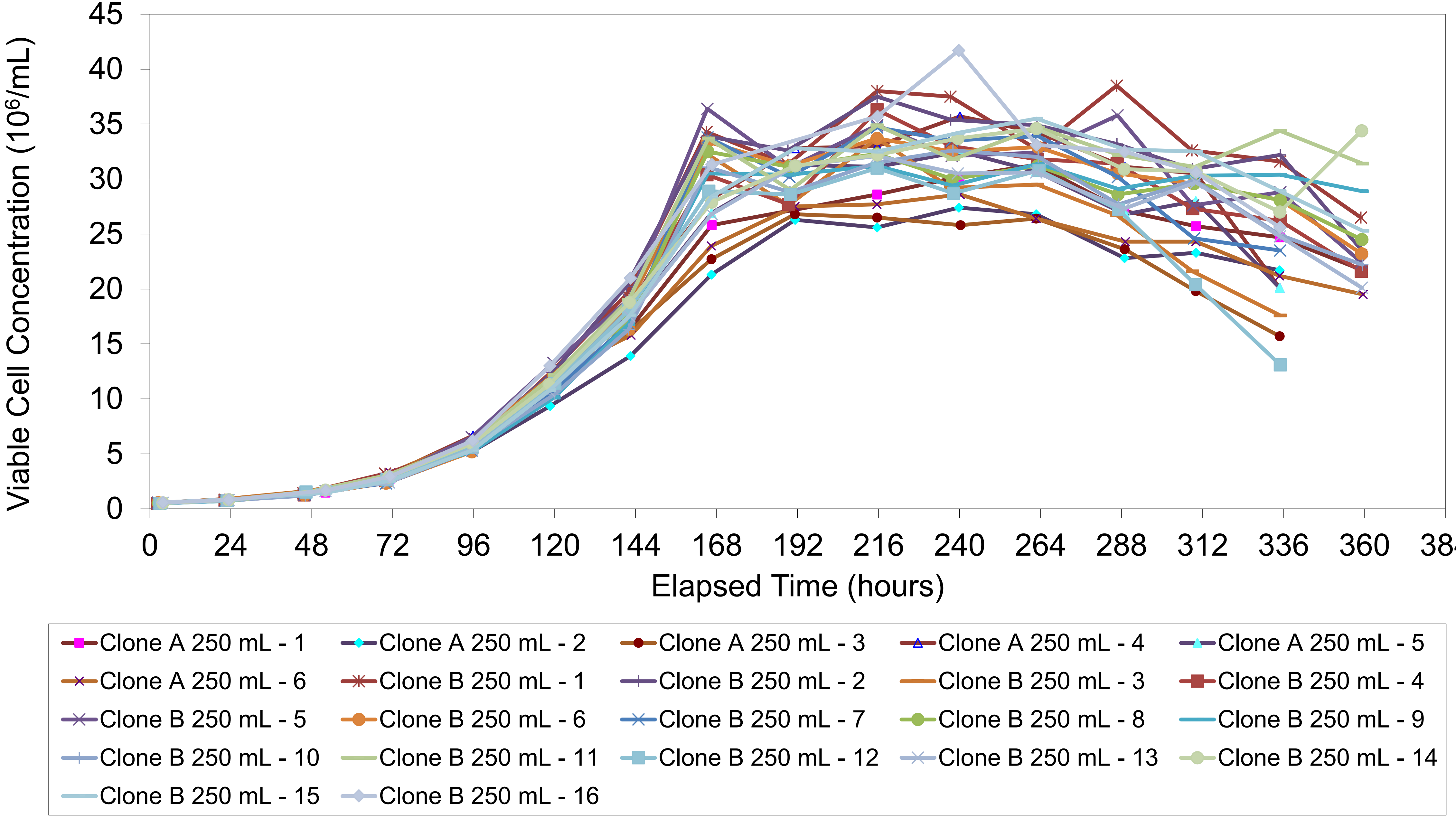

Figure 4 : Viability profile for Clone A & B across different process parameter conditions in 250 mL vessels across different culture days

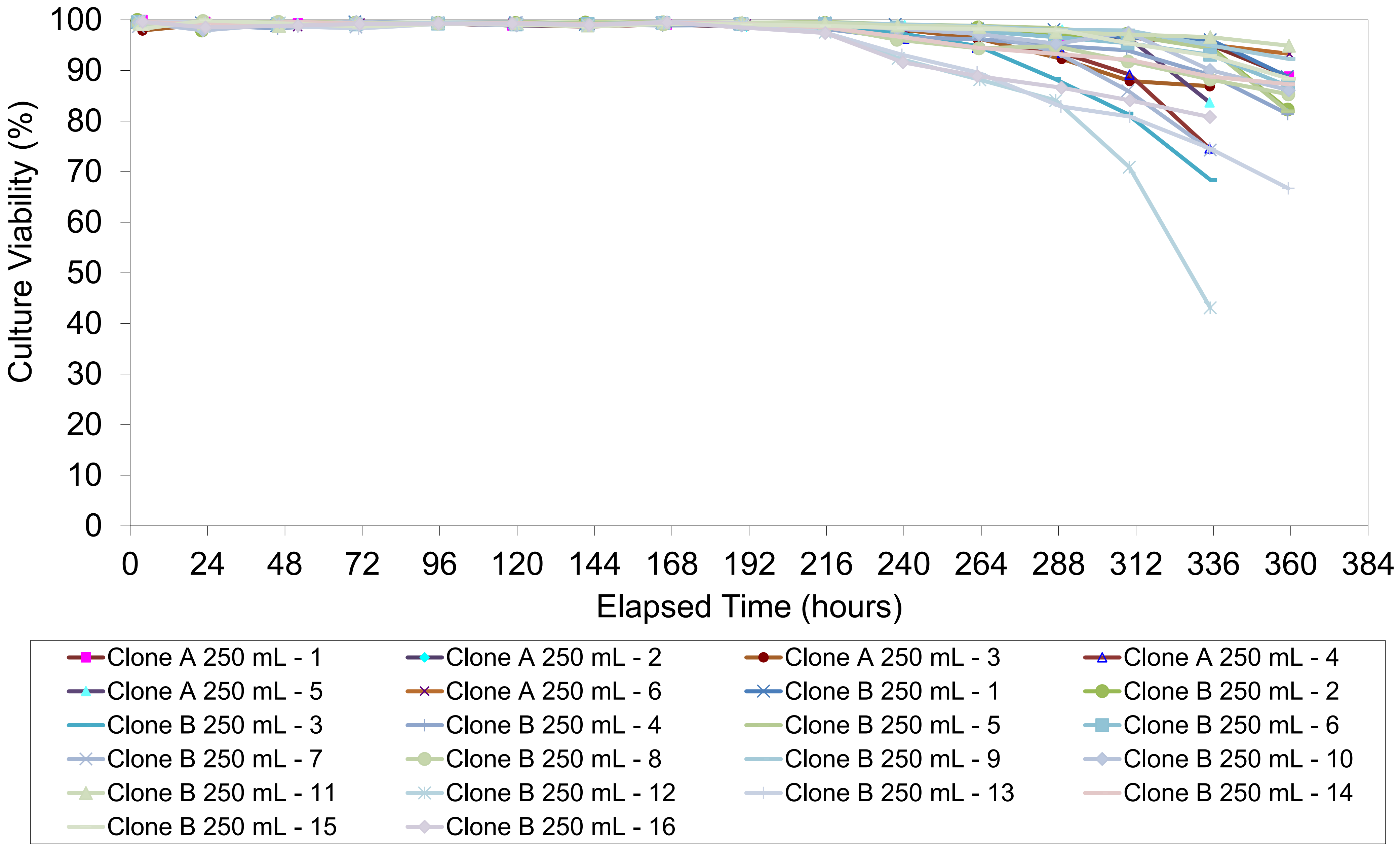

Figure X : Lactate concentration for Clone A & B across different process parameter conditions in 250 mL vessels across different culture days

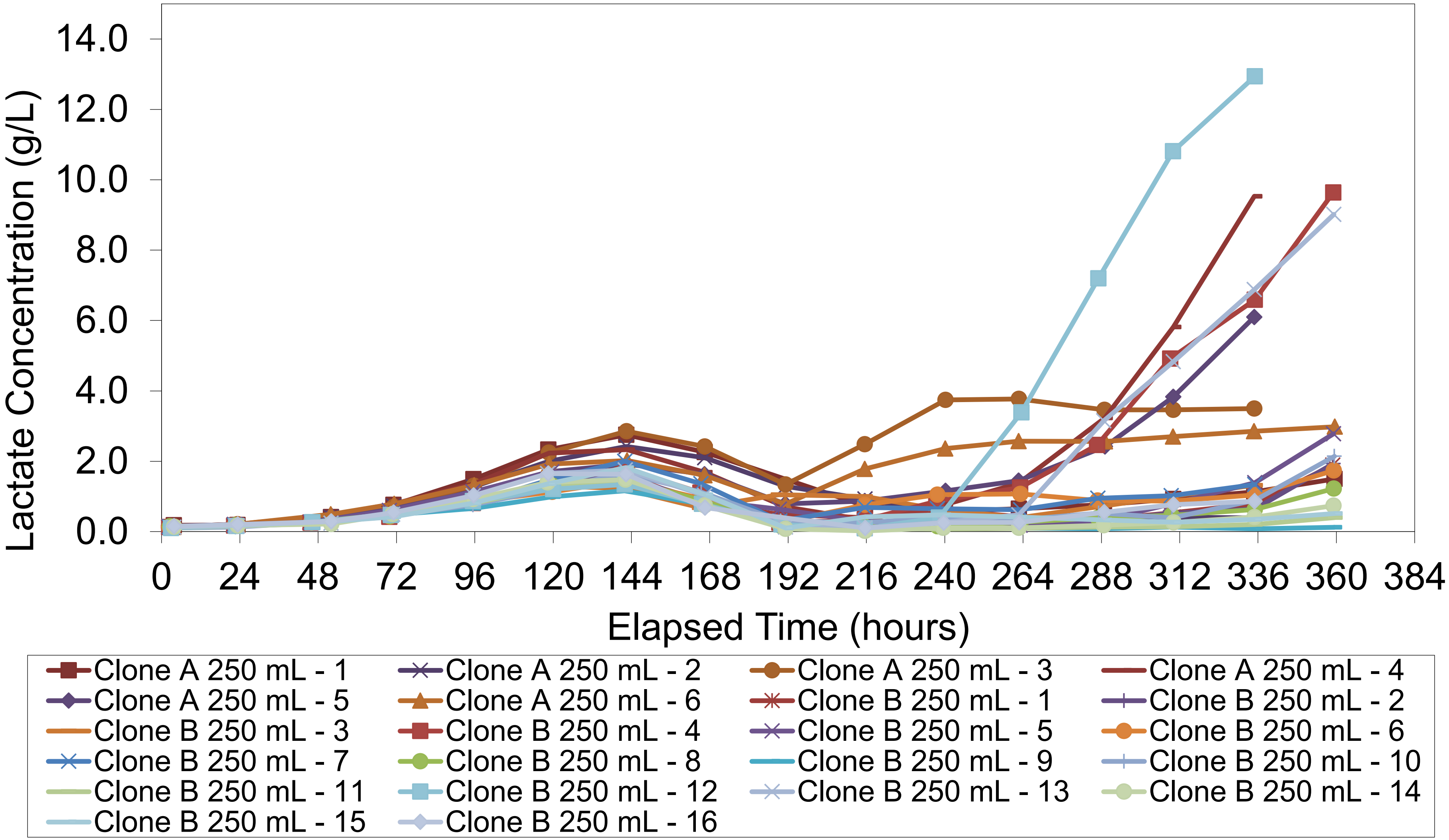

Figure X : Ammonia concentration for Clone A & B across different process parameter conditions in 250 mL vessels across different culture days

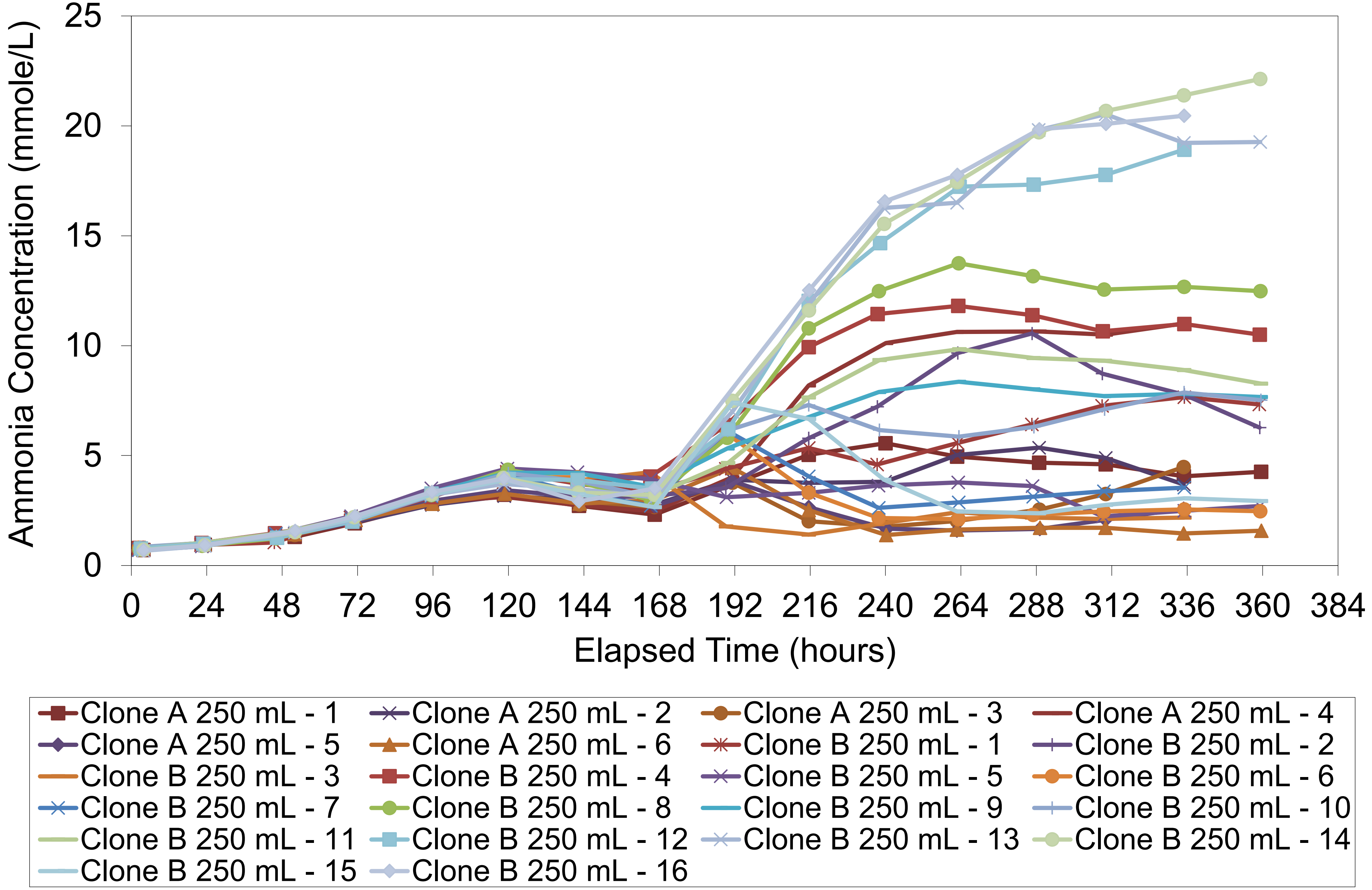

Clone B (3F7) viable cell concentration profile for Figure 5 titer plot (Single/Multi-Objective Recommendation Option 1 (No 4C2))

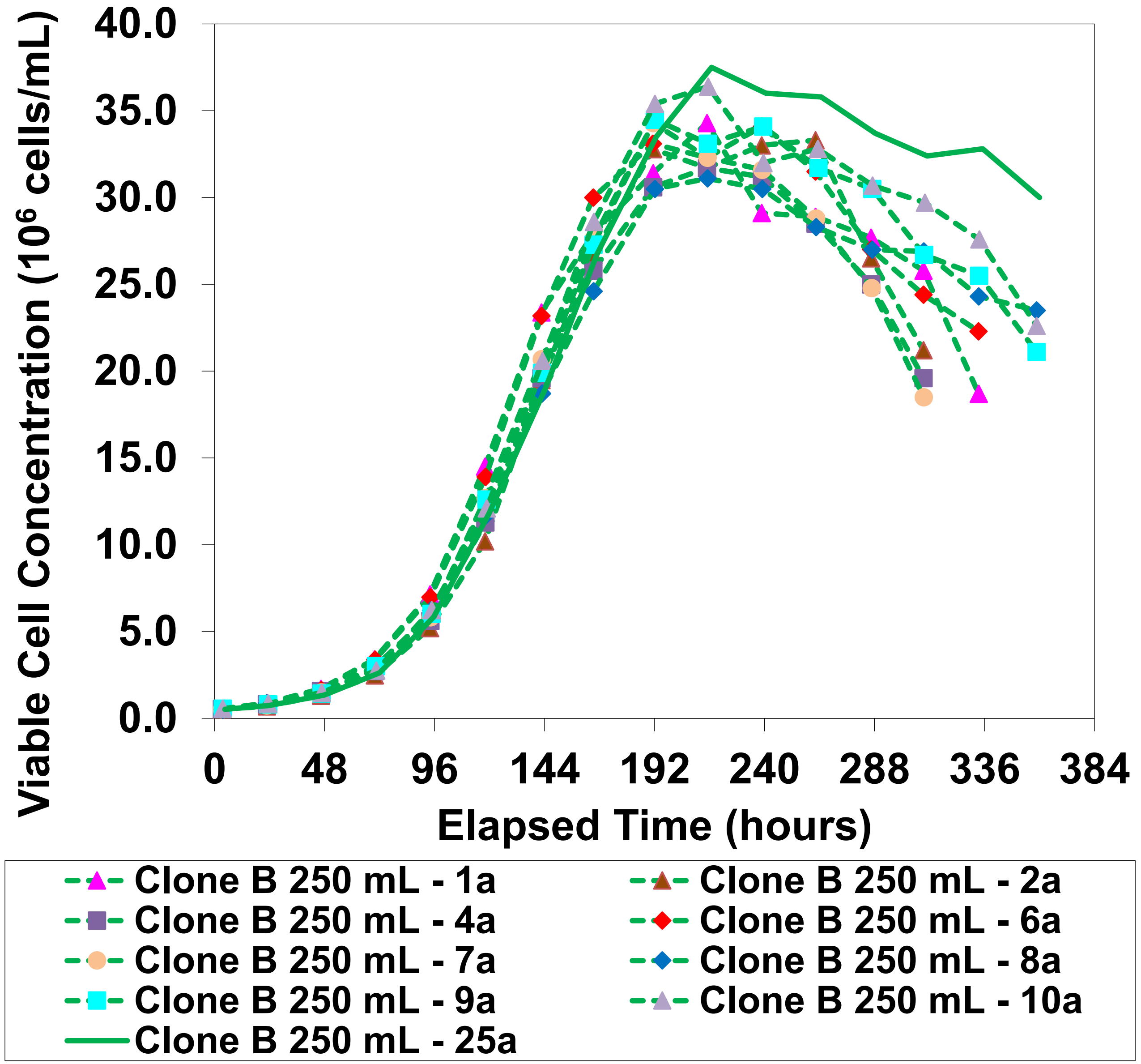


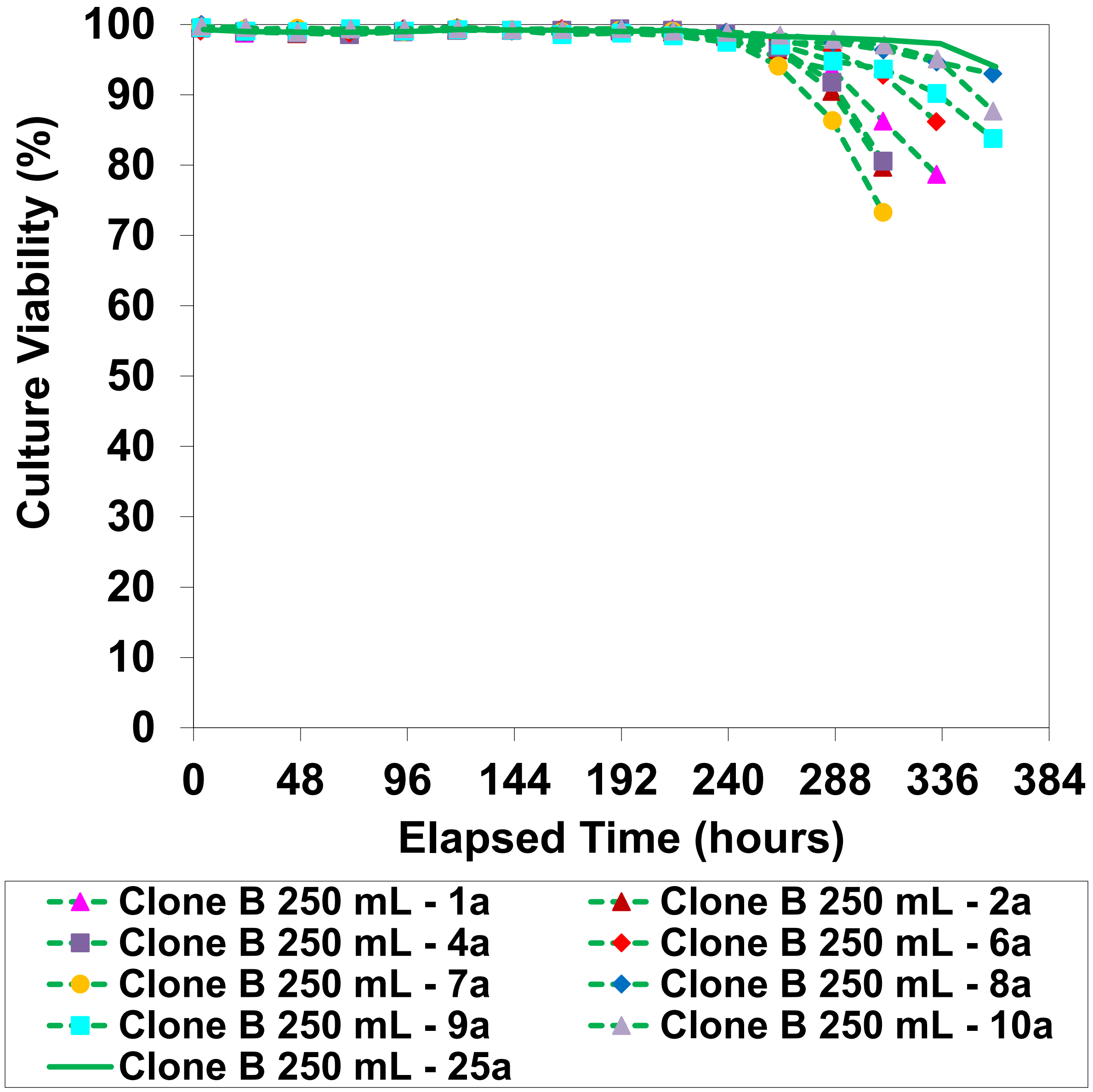

Clone B (3F7) lactate and ammonia concentration profile for Figure 5 titer plot (Single/Multi-Objective Recommendation Option 1 (No 4C2))

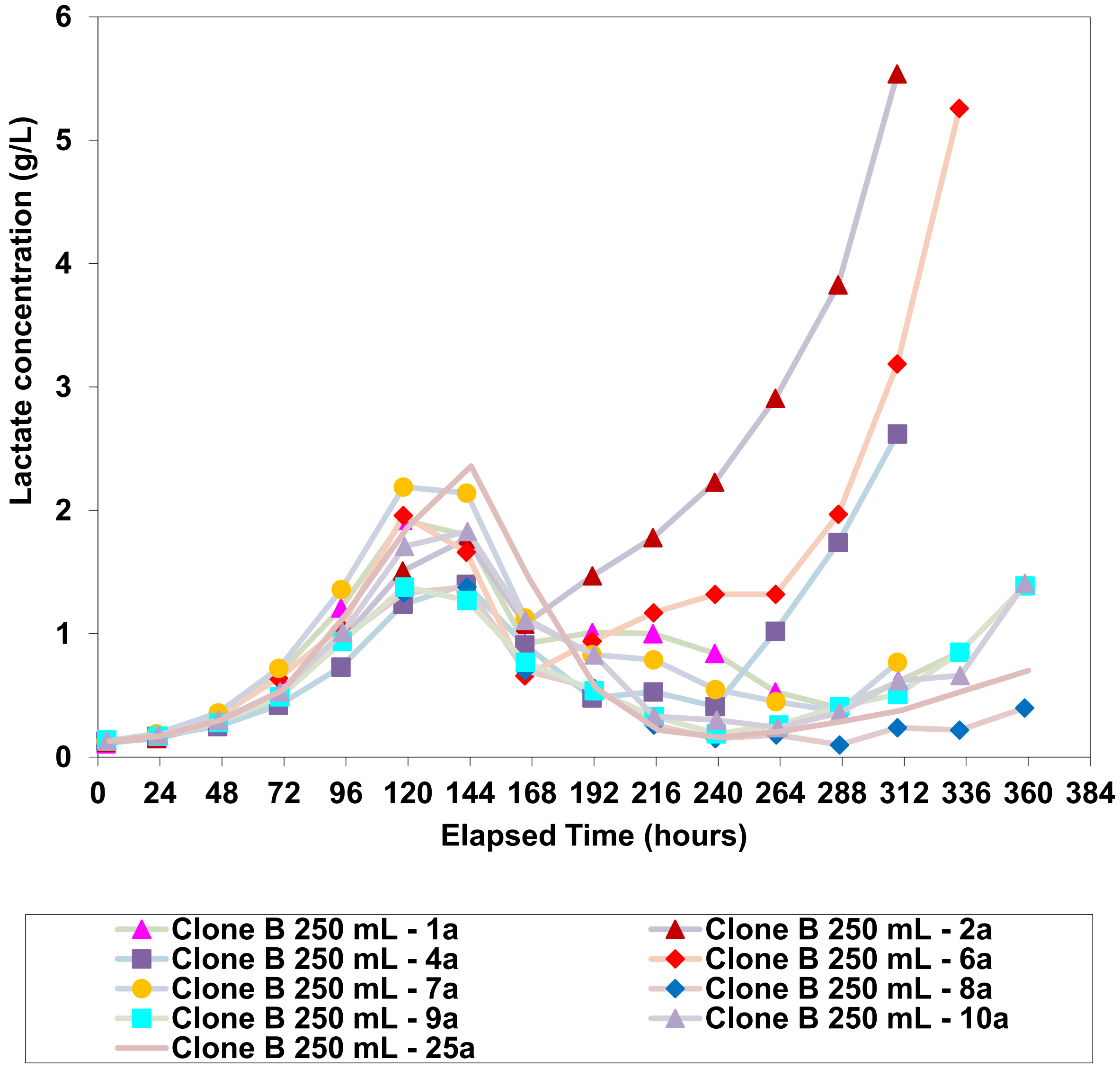


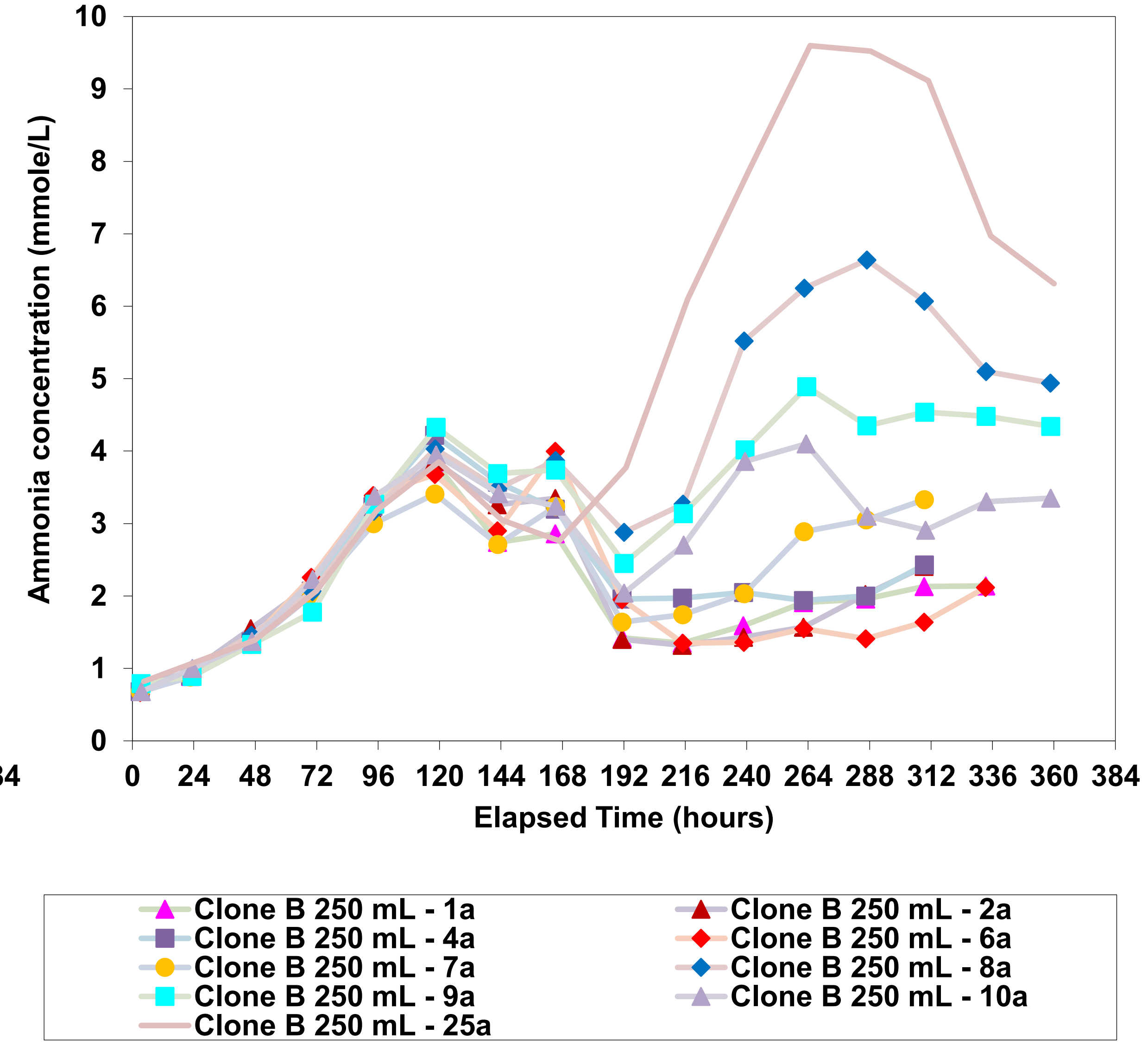

Clone B (3F7) antibody concentration profile for Figure 5 titer plot (Single/Multi-Objective Recommendation Option 1 (No 4C2))

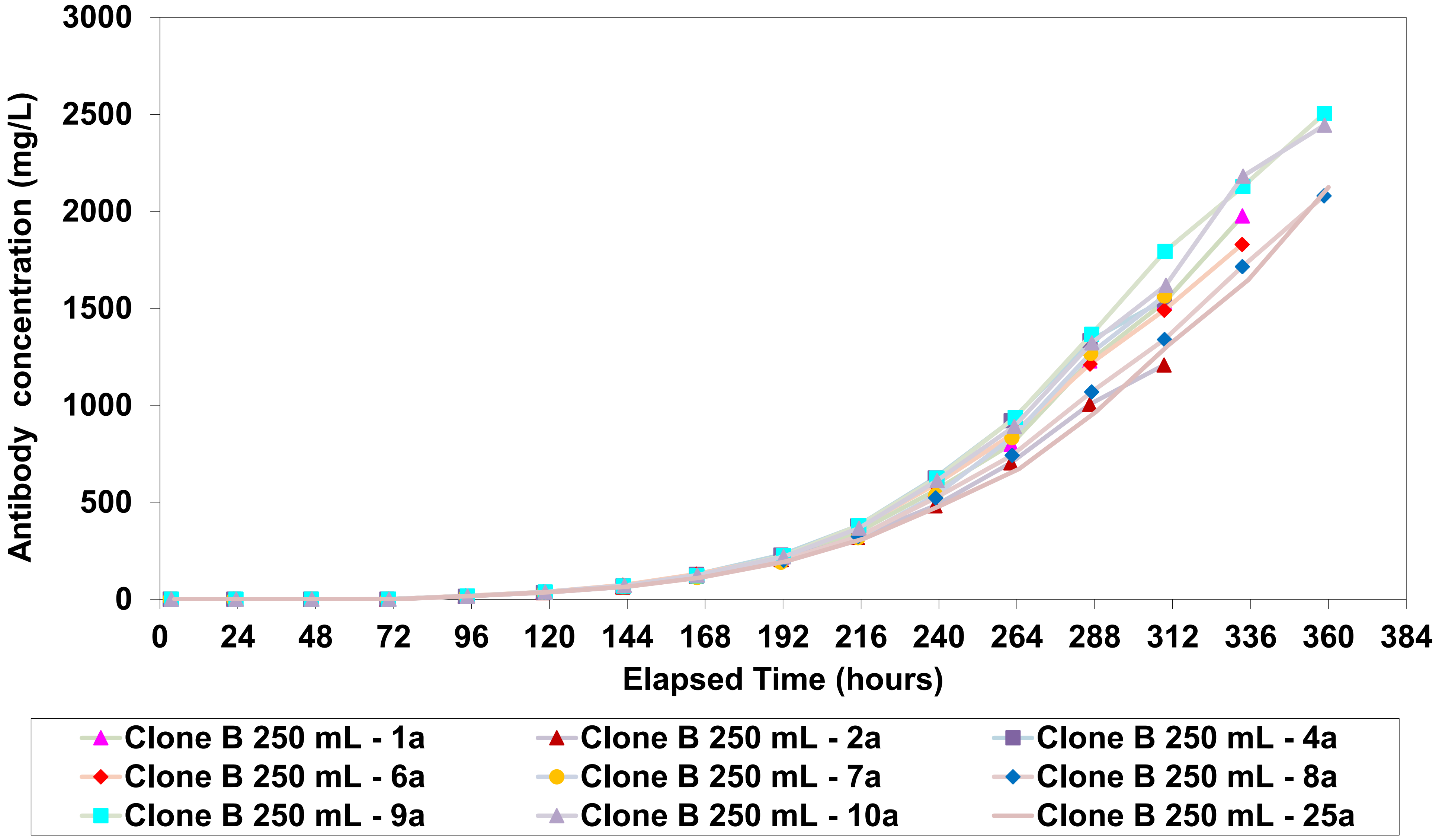

Clone A (1D11) VCC profile for Figure 5 titer plot

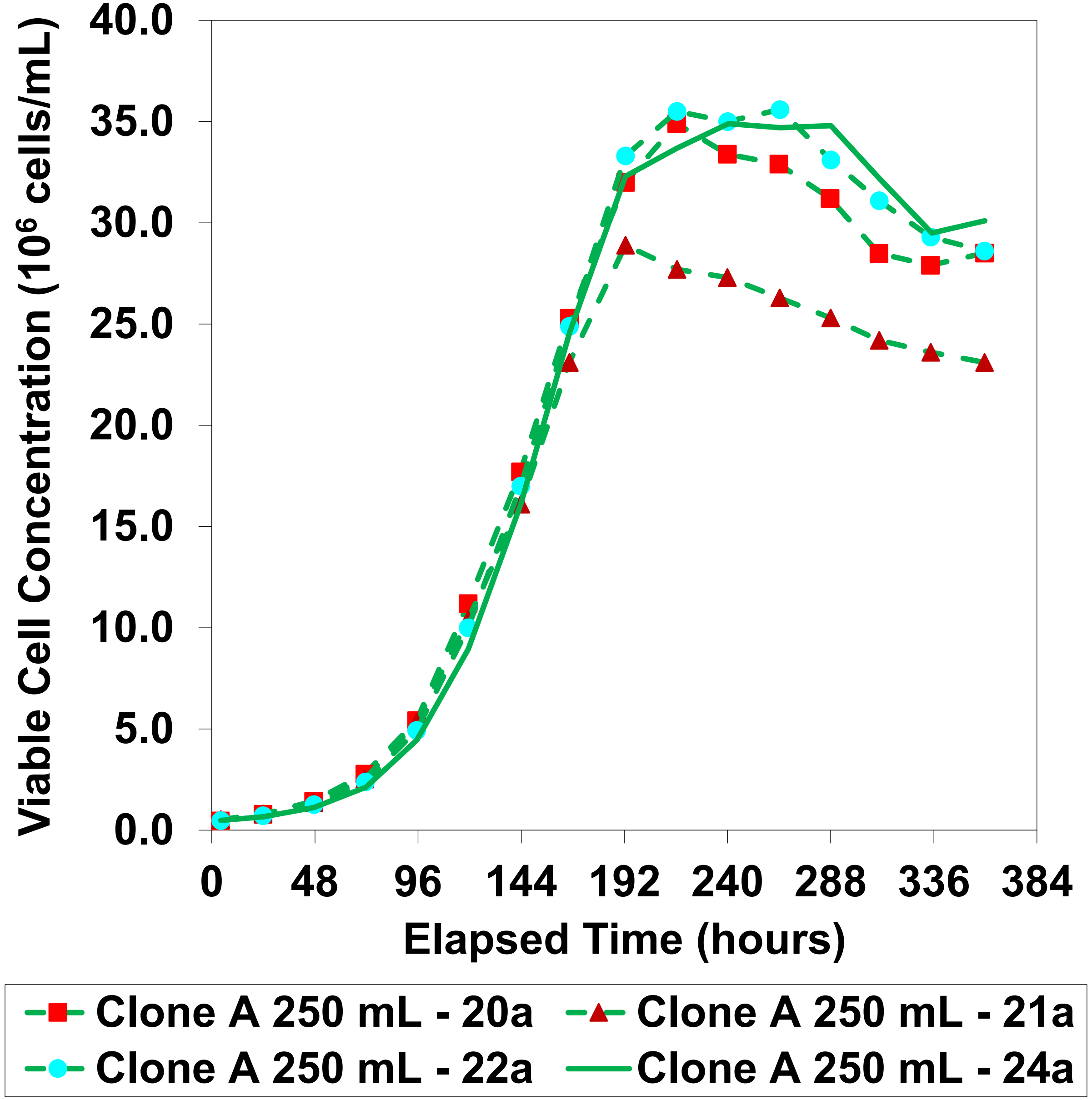


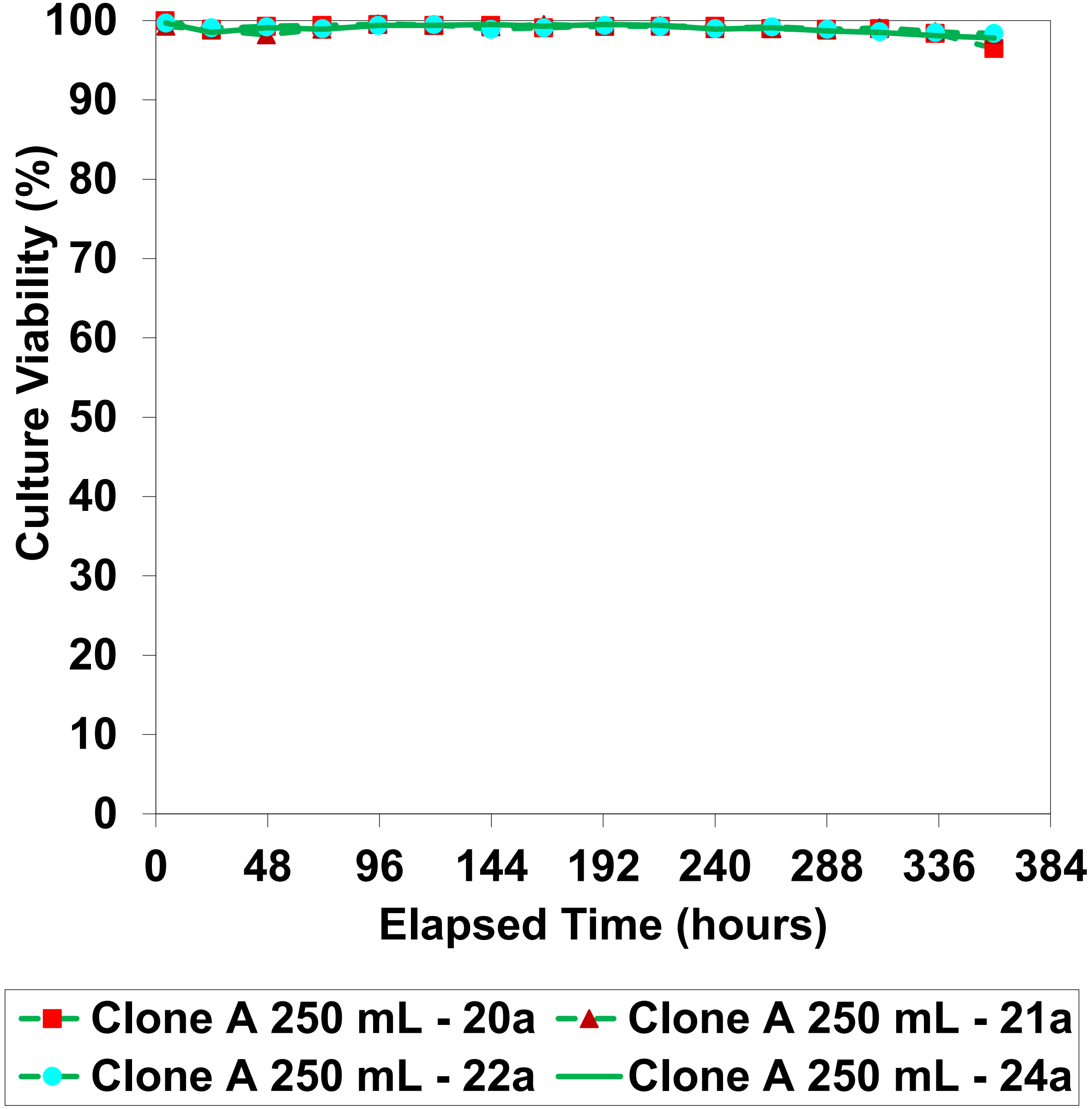

Clone A (1D11) lactate and ammonia profile for Figure 5 titer plot

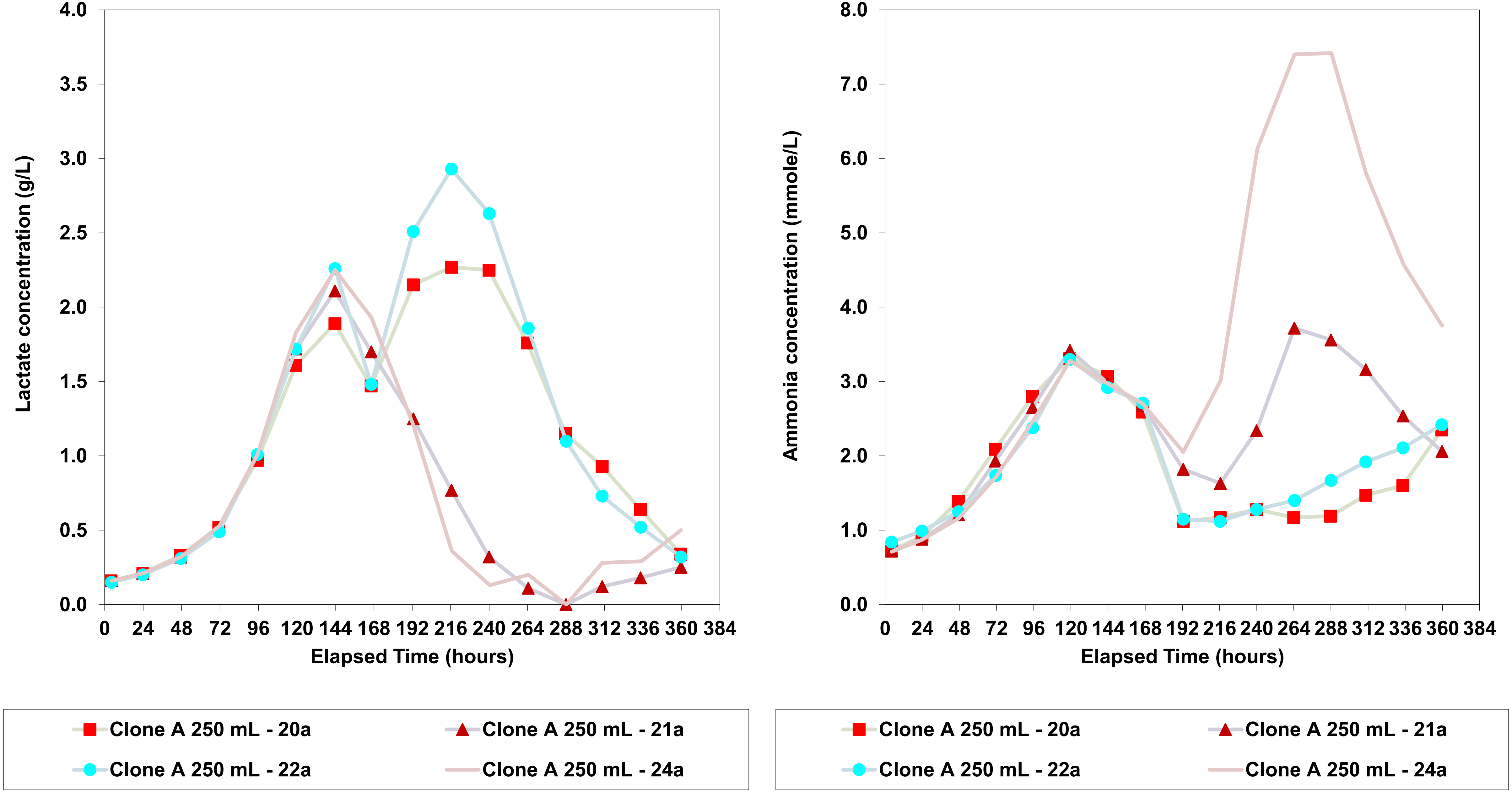

Clone A (1D11) antibody concentration profile for Figure 5 titer plot

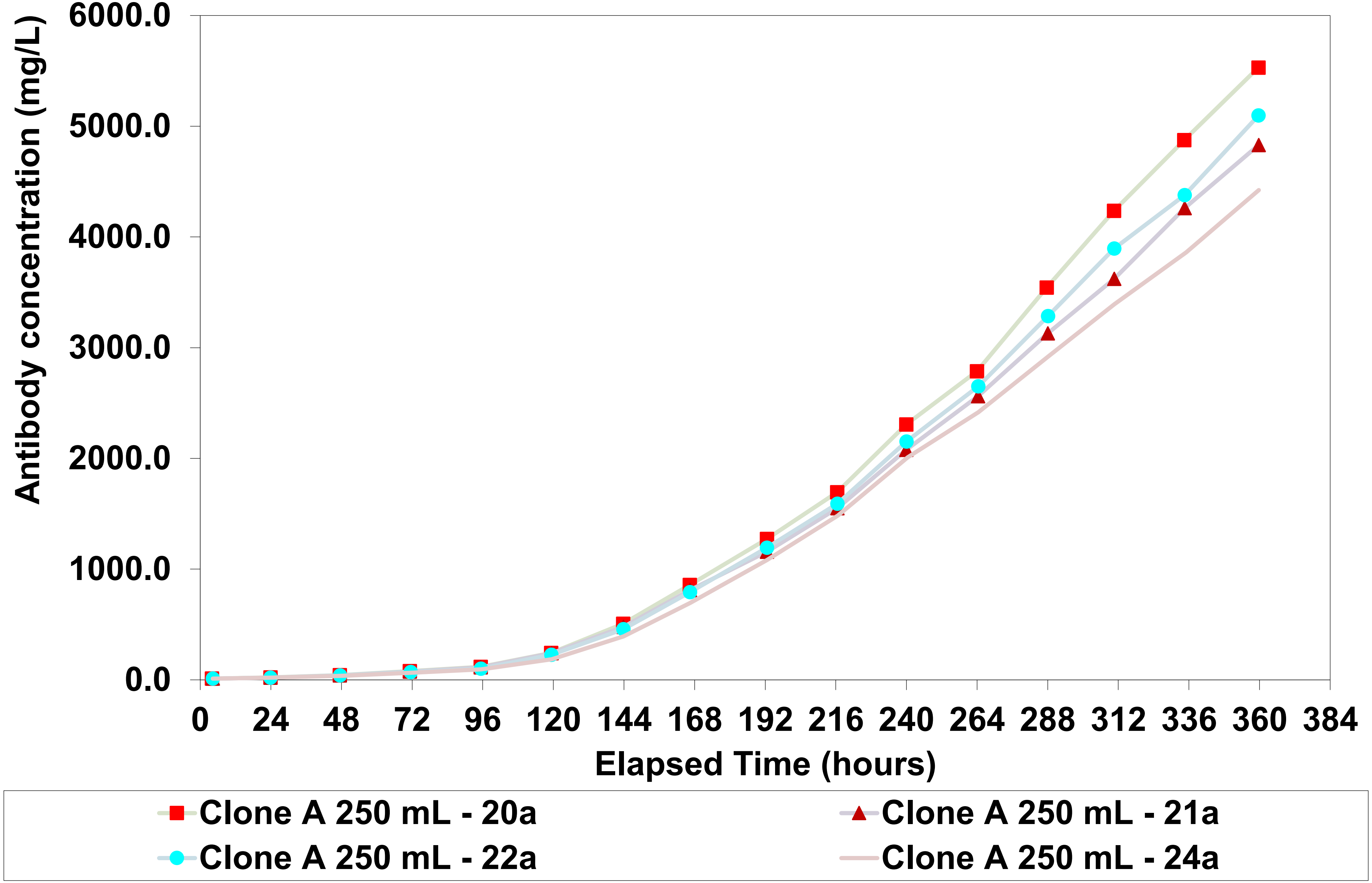

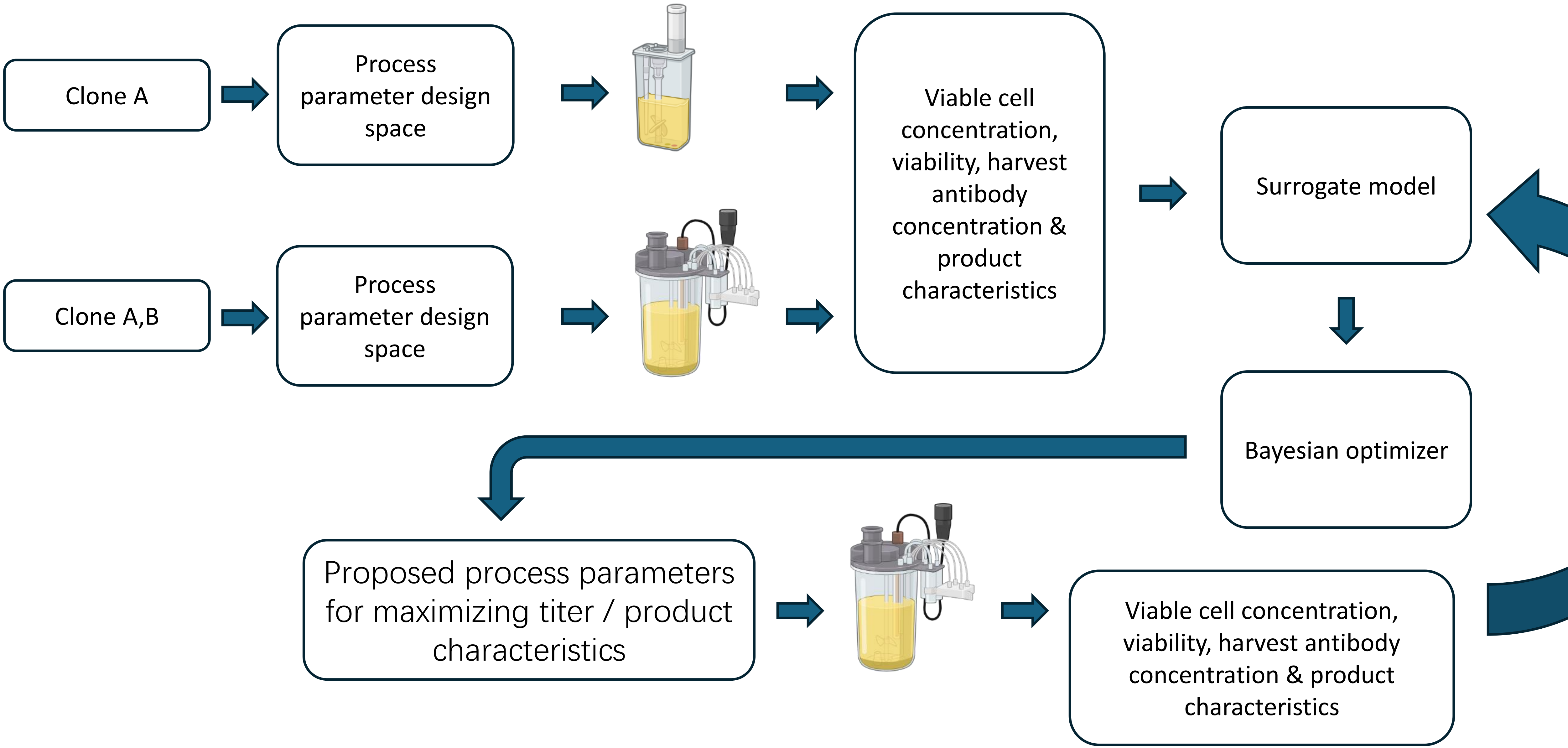
Clone A
Process parameter design space
Clone A,B
Process parameter design space
Viable cell concentration, viability, harvest antibody concentration & product characteristics
Surrogate model
Bayesian optimizer
Proposed process parameters for maximizing titer / product characteristics
Viable cell concentration, viability, harvest antibody concentration & product characteristics
